# Structural studies suggest CCDC127 as a novel membrane contact site protein in the mitochondrial intermembrane space

**DOI:** 10.64898/2025.12.11.693622

**Authors:** Tobias Bock-Bierbaum, Karina von der Malsburg, Ashwin Karthick Natarajan, Christine Zarges, Kerem Can Akkaya, Amjad Aladawi, Katja Noll, Sibylle Jungbluth, Claudia Schirra, Ying Zhu, Nils Cremer, Carola Bernert, Fan Liu, Martin Lehmann, Jan Riemer, Martin van der Laan, Oliver Daumke

**Affiliations:** Structural Biology, Max-Delbrück Center for Molecular Medicine in the Helmholtz Association, Robert-Rössle-Straße 10, 13125 Berlin, Germany; Medical Biochemistry and Molecular Biology, Center for Molecular Signaling, PZMS, Pharma Science Hub, University of Saarland, Homburg, Germany; 3Redox Metabolism Group, Institute for Biochemistry, and Cologne Excellence Cluster on Cellular Stress Responses in Aging-Associated Diseases (CECAD), University of Cologne, 50674 Cologne, Germany; Leibniz-Institut für Molekulare Pharmakologie (FMP), Robert-Rössle-Straβe 10, 13125 Berlin, Germany; Cellular Neurophysiology, Center for Integrative Physiology and Molecular Medicine (CIPMM), University of Saarland, 66421 Homburg, Germany; 6Institute of Chemistry and Biochemistry, Freie Universität Berlin, Takustraße 6, 14195 Berlin, Germany

## Abstract

Mitochondria feature a sophisticated membrane architecture, with a planar mitochondrial outer membrane (MOM) and a folded inner membrane (MIM). Due to the remarkable adaptability of mitochondria, a proteinaceous network in the intermembrane space (IMS) was proposed to confer both stability and flexibility. However, components of such scaffolds, tentatively termed the ’mitoskeleton’, have remained largely elusive. The mitochondrial contact site and organizing system (MICOS), a central organizer of mitochondrial membrane architecture, was suggested to participate in ’mitoskeleton’ formation. Here, we structurally characterize the coiled-coil domain-containing 127 (CCDC127) protein, a putative interactor of MICOS. We show that CCDC127’s amino-terminal transmembrane region is anchored in the MOM and the bulk soluble part exposed to the IMS. A crystal structure of CCDC127’s central coiled-coil displays a parallel dimer which further oligomerizes into tetramers. We demonstrate that the carboxy-terminal helical bundle (CHB) domain dimerizes to create a peripheral membrane-binding site. Supported by electron microscopy data, we propose a structural model of CCDC127 as intramitochondrial membrane contact site protein mediating the structural organization of the IMS as part of the ’mitoskeleton’.

## INTRODUCTION

Mitochondria are probably the most flexible and versatile organelles within eukaryotic cells constantly adapting their morphology and metabolic performance according to the needs of the cell. Their filigree inner architecture is characterized by an enveloping mitochondrial outer membrane (MOM) handling communication with other organelles and the cytosol together with a mitochondrial inner membrane (MIM) that folds up into cristae membranes harboring the oxidative phosphorylation machinery. The biogenesis and maintenance of such complex membrane structures requires a plethora of assembly factors, chaperones and scaffolding proteins. In addition, mitochondria are constantly exposed to incoming and outgoing signals triggering fusion and fission events, adaptive ultrastructural remodeling and biochemical re-programming (Bennett *et al*, 2022; Chen *et al*, 2025; Cheng *et al*, 2023; Daumke & van der Laan, 2025; Giacomello *et al*, 2020; Monzel *et al*, 2023; Pernas & Scorrano, 2016; Pfanner *et al*, 2025; Quintana-Cabrera & Scorrano, 2023; Spinelli & Haigis, 2018; Suomalainen & Nunnari, 2024). Surrounded by the MOM and MIM, the narrow intermembrane space (IMS) appears to be a particularly challenged and vulnerable compartment (Edwards *et al*, 2020; Hohorst *et al*, 2025; Weith *et al*, 2025). Across an extensive mitochondrial surface area, MIM and MOM are closely apposed, with a constant small diameter of the IMS in the range of 20-50 nm. At crista junctions, cristae membranes originate to form extended tubular or lamellar invaginations. These regions of exceptionally high membrane curvature are stabilized by the mitochondrial contact site and cristae organizing system (MICOS), a large multi-subunit membrane scaffold with at least two membrane-bending protein components, Mic10 and Mic60 (Barbot *et al*, 2015; Bock-Bierbaum *et al*, 2022; Bohnert *et al*, 2015; Daumke & van der Laan, 2025; Eramo *et al*, 2020; Guarani *et al*, 2015; Hessenberger *et al*, 2017; Klecker & Westermann, 2020; Kondadi & Reichert, 2024; Stephan *et al*, 2020; Tarasenko *et al*, 2017). Moreover, MICOS forms physical contact sites between MIM and MOM through protein-protein interactions with the sorting and assembly machinery (SAM complex) and the general preprotein translocase of the MOM, the TOM complex (Harner *et al*, 2011; Hoppins *et al*, 2011; Huynen *et al*, 2016; Ott *et al*, 2012; Sastri *et al*, 2017; Tang *et al*, 2020; von der Malsburg *et al*, 2011). Such intra-mitochondrial membrane contact sites appear to be essential for IMS architecture. In MICOS-deficient mitochondria, cristae membranes are detached from the MIM and accumulate as stacked membrane sheets within the matrix, separating the intracristal space from the boundary IMS (Colina-Tenorio *et al*, 2020; Daumke & van der Laan, 2025; Kondadi & Reichert, 2024; Mukherjee *et al*, 2021).

Besides the critical role of MICOS, only little is known about factors that assure the structural and functional integrity of the narrow IMS compartment. MICOS complexes were shown to form helical punctate patterns that wind around tubular mitochondrial segments (Hoppins *et al*., 2011; Itoh *et al*, 2013; Stoldt *et al*, 2019). These peculiar arrangements are thought be part of a hypothetical IMS scaffolding machinery that has been address occasionally as the ‘skeletal structure’ or ‘mitoskeleton’, despite the fact that its composite molecular architecture is largely unknown (Colina-Tenorio *et al*., 2020; Hobbs *et al*, 2001; Hoppins *et al*., 2011; Ishihara *et al*, 2022; Stoldt *et al*., 2019). The first discovered direct membrane contact sites formed across the mitochondrial envelope were transient super-complex assemblies of the TOM complex and the preprotein translocase of the inner membrane, TIM23, holding a two-membrane spanning precursor protein *en route* to the mitochondrial matrix (Chacinska *et al*, 2010; Gomkale *et al*, 2021; Reichert & Neupert, 2002; Wang & Nussberger, 2024; Yang *et al*, 2025; Zhou *et al*, 2023). Physical interactions between the fusion machineries of MOM and MIM and a putative two-membrane-spanning mtDNA segregation machinery were also reported (Boldogh *et al*, 2003; Hobbs *et al*., 2001; Sesaki & Jensen, 2004; Wong *et al*, 2003). However, only the discovery of the mitochondrial intermembrane space bridging (MIB) complex formed by MICOS together with the sorting and assembly machinery (SAM) of the MOM revealed a stable, yet flexible structure that may be able to withstand and cushion the lateral and transversal pressures exerted on MOM and MIM during mitochondrial fusion and fission, contact site formation and transport along cytoskeletal elements (Colina-Tenorio *et al*., 2020; Daumke & van der Laan, 2025; Hoppins & Nunnari, 2012; Kozjak-Pavlovic, 2017; Ott *et al*., 2012; Schorr & van der Laan, 2018; Stoldt *et al*., 2019; Tang *et al*., 2020).

Despite substantial progress in understanding mitochondrial dynamics and internal membrane architecture, our knowledge about the molecular identity and structural organization of mitochondria-shaping and -scaffolding proteins remains limited. Identification of candidate proteins appears complicated due to potential redundancy and adaptive processes in classical knock-out mutant cells (Zarges *et al*, 2025a). A recent study described the two-membrane-spanning AAA^+^ ATPase ATAD3A protein as a potential component of a proteinaceous meshwork that connects MIM and MOM and may control the regular structure of the IMS, thereby influencing both its mechanical and signaling properties (Arguello *et al*, 2021). Such central coordinating functions may explain the pleiotropic phenotypes of ATAD3A mutant cells (Chen *et al*, 2023).

We aimed to identify potential novel components that may structurally support or organize the IMS through MOM and MIM scaffolding or tethering based on their structural features. The prototypical IMS-organizing protein subunit of the MICOS complex, Mic60, is anchored to the MIM via a transmembrane segment (TMS) and combines extended coiled-coil regions with helical bundle domains to oligomerize into a scaffold and bind to membrane surfaces (Bock-Bierbaum *et al*., 2022; Hessenberger *et al*., 2017). Of note, a recent study noticed a protein termed CCDC127 (for <u>C</u>oiled<u>-C</u>oil <u>D</u>omain <u>C</u>ontaining protein 127) which can be chemically crosslinked to the human MICOS components MIC19, MIC25 and MIC60 in mitochondria (Fig. 1A) (Zhu *et al*, 2024). CCDC127 homologues are found in vertebrates, but are not present in insect, fungal and plant species. Earlier studies had identified CCDC127 as a transcription factor regulating the expression of *hsp70* genes in a yeast-two-hybrid screen (Saito *et al*, 2016) or as a mitochondrial surface-exposed protein that impacts lipid shuttling between mitochondria and lipid droplets (Xia *et al*, 2023).

**Figure 1:**
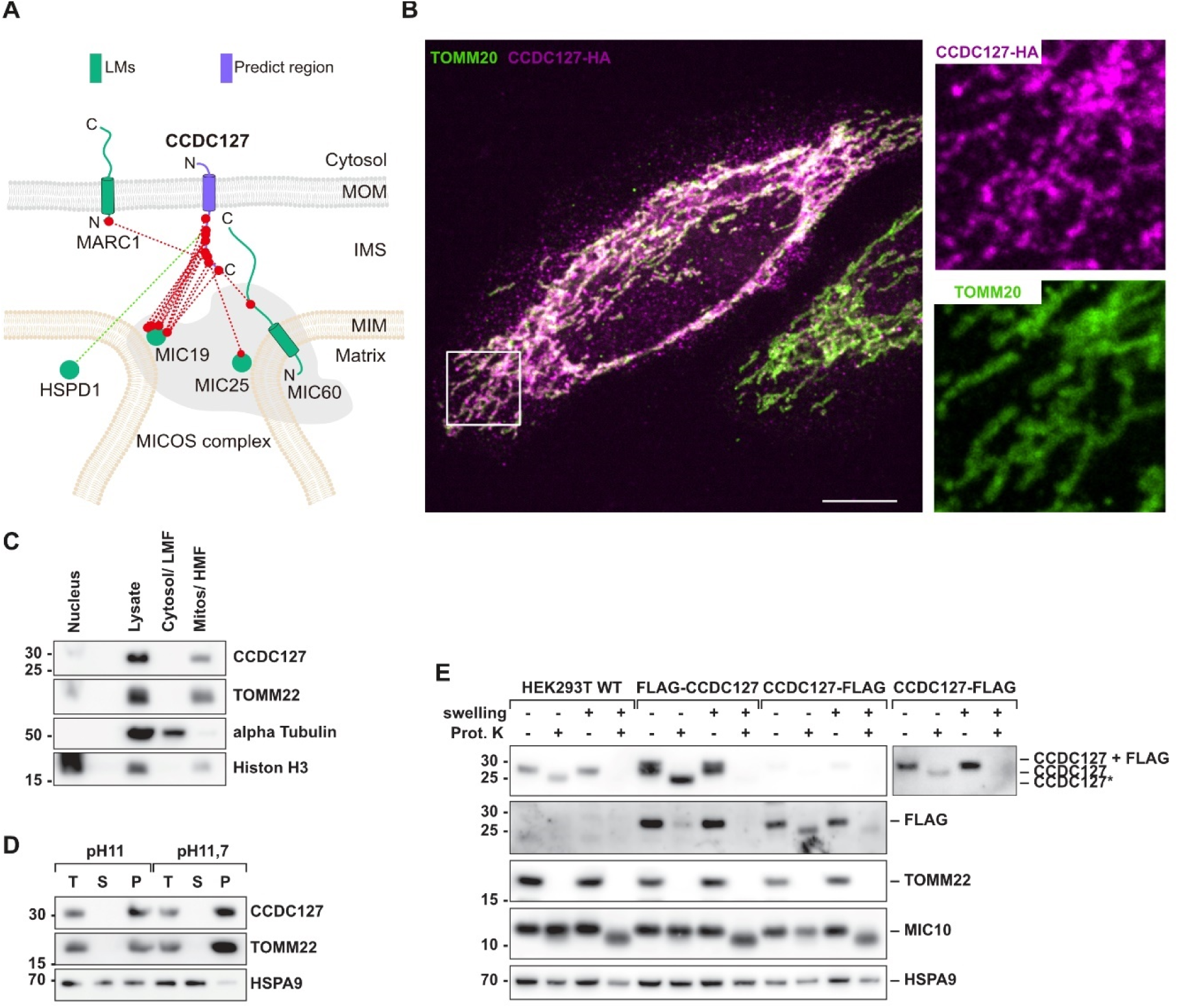
CCDC127 is a mitochondrial outer membrane protein exposed to the inter-membrane space. **A.** Cross-link map of CCDC127 and its interacting proteins derived from two XL-MS datasets. Localization markers (LMs) are shown in green; CCDC127 is shown in purple. The cross-links are shown in red. The cross-linking assisted spatial proteomics (CLASP) annotation of CCDC127 is supported by direct connections to two IMS LMs, one IMM LM and one MOM LM. “Predict region” indicates the protein, for which a CLASP prediction was made (Zhu *et al*., 2024). **B.** Confocal dual-color microscopy of CCDC127-HA and TOMM20 in PFA-fixed Hela cells. Scale bar and inserts: 15 μm. The right upper insert shows CCDC127-HA and the lower insert TOMM20. **C.** Cellular fractionation experiment showing the subcellular localization of CCDC127. LMF, light membrane fraction; HMF, heavy membrane fraction. **D.** Alkaline extraction of isolated mitochondria demonstrating the membrane association of CCDC127. T, total; P, pellet; S, supernatant. **E.** Swelling and proteinase K accessibility assay of HEK293T wild-type, HEK FLAG-CCDC127 and HEK CCDC127-FLAG mitochondria, indicating the N-out topology of CCDC127. Top right panel showing a longer exposure of the CCDC127 western blot for CCDC127-FLAG mitochondria.

Here, we describe CCDC127 as a previously uncharacterized mitochondrial MOM protein that is exposed to the IMS and likely cooperates with MICOS. Structural and biochemical studies revealed a dimeric architecture of CCDC127 characterized by an N-terminal TMS that anchors the protein to the MOM, a central coiled-coil domain dimer, which may span across the IMS, and a C-terminal helical bundle domain, which acts as a peripheral membrane-binding site. Our data suggest that CCDC127 bridges MOM and MIM to function as a novel membrane contact site protein that stabilizes mitochondrial membrane architecture.

## RESULTS

### CCDC127 is an integral MOM protein exposed to the IMS

We carefully re-examined the subcellular localization and membrane topology of CCDC127 in HEK293T cells. Confocal dual-color fluorescence microscopy of endogenous antibody-labelled CCDC127 (Fig. S1A) or transiently over-expressed CCDC127 containing a C-terminal HA-tag (Fig. 1B) showed a clear co-localization with the MOM marker protein TOMM20 in mitochondria, but their spreading throughout the mitochondrial network was clearly distinct. While TOMM20 showed a uniform distribution, CCDC127 was found in a punctate pattern. Biochemical fractionation of wild-type HEK293T cells followed by immunoblot analysis confirmed mitochondrial localization of CCDC127 (Fig. 1C). Alkaline extraction experiments using isolated mitochondria demonstrated that CCDC127 behaves like a canonical integral membrane protein, because it was found in the pellet fraction after treatment at pH 11 and pH 11.7 and ultracentrifugation (Fig. 1D).

To determine the sub-mitochondrial localization and transmembrane topology of CCDC127, we generated variants of the protein carrying a FLAG-tag either at the N- or at the C-terminus. We isolated mitochondria from wild-type, FLAG-CCDC127 and CCDC127-FLAG cells and performed a protease accessibility assay. Mitochondria were either kept under iso-osmotic conditions or subjected to a hypo-osmotic shock (swelling) prior to proteinase K treatment. Hypo-osmotic swelling of mitochondria leads to rupture of the MOM rendering IMS-exposed proteins accessible to proteolytic degradation. Under iso-osmotic control conditions, only protein domains exposed to the cytosolic surface of the MOM are removed by the protease. Untagged CCDC127 detected with a specific antibody migrated slightly faster in SDS-PAGE when intact mitochondria treated with proteinase K were analyzed indicating that a small protein fragment was degraded while the bulk of the protein remained protected by the MOM (Fig. 1E). Similar patterns were observed with both FLAG-tagged CCDC127 variants. However, when an anti-FLAG antibody was used for visualization, only the variant tagged at the C-terminus could still be detected after protease treatment, whereas the N-terminally tagged variant was not observed (Fig. 1E). Upon hypo-osmotic swelling of mitochondria, all CCDC127 versions and their respective epitopes became accessible to proteinase K, comparable to the IMS-exposed MIM-protein MIC10. Matrix-localized HSPA9 remained protease-resistant under all conditions (Fig 1E). We conclude that CCDC127 is an integral MOM protein that exposes a short N-terminal region to the cytosol, whereas the remainder of the protein localizes to the IMS. Localization and topology agree with a putative role of CCDC127 in the spatial organization of the IMS.

### A parallel coiled-coil mediates CCDC127 dimerization and oligomerization

To obtain a functional understanding of CCDC127, we aimed for the structural characterization of the protein. While full-length CCDC127 remained inaccessible for structural studies (see below), we recombinantly expressed and purified a CCDC127-derived construct comprising the predicted coiled-coil domain (CCDC127^CC^, residues 48-120) (Fig. 2A, Fig. S1B). Crystals of this coiled-coil construct were obtained in space group P3_2_21 and diffracted to 2.45 Å resolution (Table S1, S2). They contained six protein copies in the asymmetric unit and displayed pseudo-translation, which was reflected in relatively high R_work_/R_free_ factors during the refinement (Table S2). The six CCDC127 molecules in the asymmetric unit formed three parallel dimeric coiled-coils. Two of these dimers further assembled into a two-fold symmetric tetramer, whereas the third dimer formed an analogues tetramer with a symmetry-related dimer from an adjacent asymmetric unit (Fig. 2B). Notably, the tetramerization interface is evolutionary conserved (Fig. 2C, Fig. S4), pointing to a functional relevance. It features salt bridges between D69 and K73 and a hydrophobic core, formed by L70 and Y74 (Fig. 2B, boxes 1 and 2).

**Figure 2:**
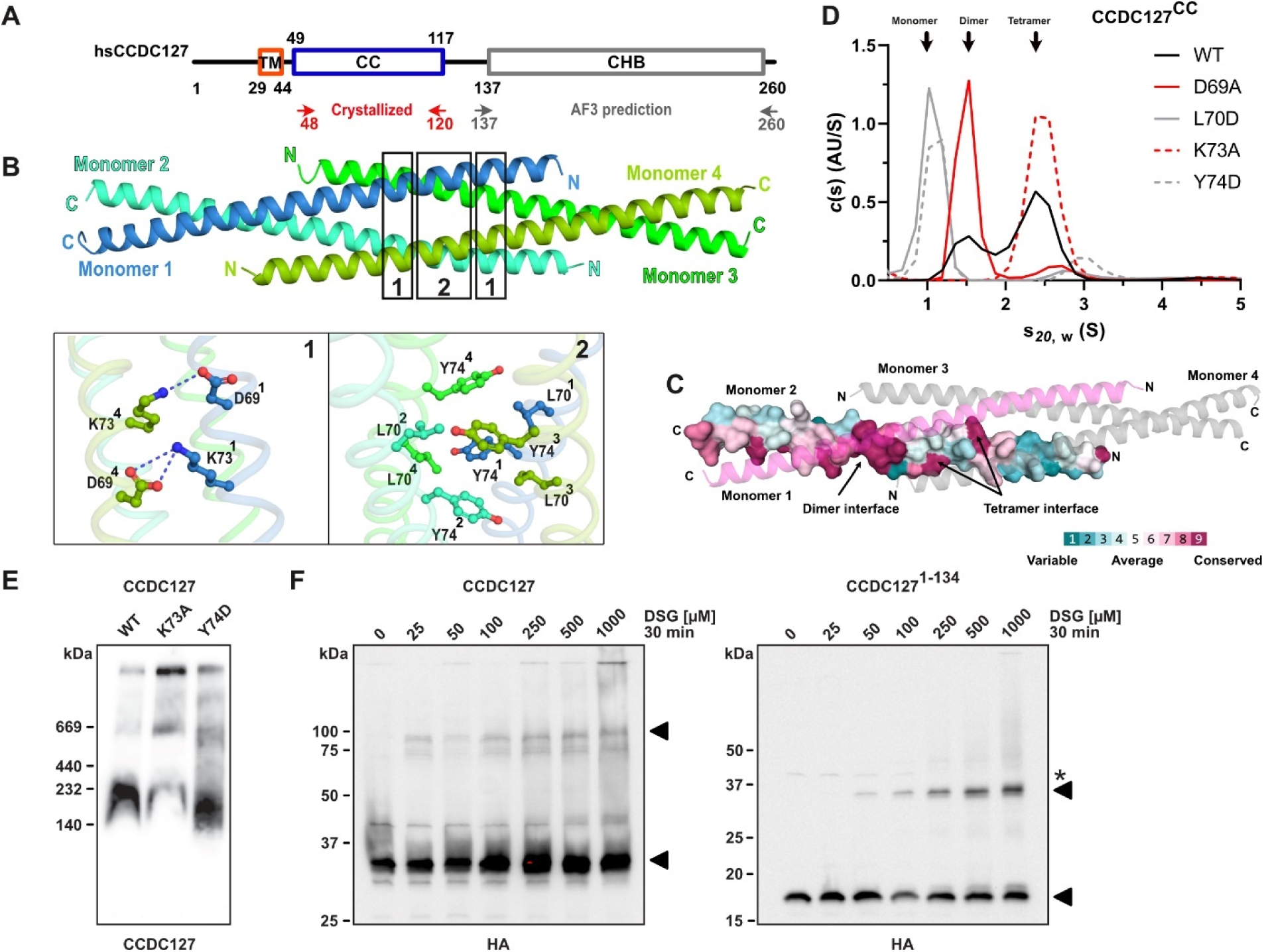
A parallel coiled-coil domain mediates dimerization and higher-order oligomerization of CCDC127. **A.** Domain architecture of human CCDC127. Structurally characterized regions are indicated. **B.** Crystal structure of the tetrameric coiled-coil domain of human CCDC127 (CCDC127^CC^, residue 48-120). Two magnified views of the tetrameric interface characterized by hydrophilic (**Box 1**) and hydrophobic (**Box 2**) patches. Colors and labels are identical. **C.** Surface conservation plot of the coiled-coil domain of CCDC127. For one monomer, each residue is colored according to its conservation and shown as surface representation. The second monomer involved in dimerization is colored in magenta and shown as cartoon representation. Monomer 3 and 4 are colored in grey. Highly conserved residues are colored magenta, variable resides in dark cyan. **D.** Sedimentation velocity measurements to determine the oligomeric state in solution of WT and variant (D69A, L70D, K73A and Y74D) CCDC127^CC^. **E.** CCDC127 protein complex analysis by BN-PAGE using detergent-solubilized isolated mitochondria from HEK293T WT cells and cells expressing the CCDC127 variants K73A and Y74D. **F.** CCDC127 WT-HA and CCDC127^1-134^-HA can be crosslinked in cells using the crosslinker DSG with increasing concentrations.

To test the functional relevance of this tetrameric interface, we introduced single amino acid alterations into the CCDC127^CC^ construct and analyzed the assembly status of the resulting coiled-coil variants by sedimentation velocity analytical ultracentrifugation (AUC, Fig. 2D) and analytical size-exclusion chromatography (SEC, Fig. S1C). In these experiments, the unmodified CCDC127^CC^ construct formed dimers and tetramers. Disruption of the central hydrophobic core at the tetramer interface through the amino acid replacements L70D or Y74D led to the disassembly of the complex into monomers. In contrast, the single D69A exchange led to the formation of a stable dimer in AUC and SEC experiments. Strikingly, the CCDC127^CC^-K73A variant almost exclusively appeared as a tetramer, most likely since the newly introduced alanine residue contributes to the hydrophobic core of the tetrameric arrangement (Fig. S1D).

To assess the assembly behavior of CCDC127 in a cellular context, we stably expressed full-length wild-type CCDC127 as well as variants carrying the amino acid alterations K73A or Y74D in CCDC127-deficient HEK293T knockout cell lines. We isolated mitochondria from these cells, solubilized them with the mild detergent digitonin and analyzed the protein extracts by blue native-PAGE (BN-PAGE). To visualize the protein complex profile of CCDC127 in the different samples, we employed immunoblotting with specific antibodies against the protein. Wild-type CCDC127 indeed migrated as a mixture of different high-molecular-weight complex forms in line with the idea that the protein oligomerizes in intact mitochondria (Fig. 2E). The K73A substitution led to the stabilization of higher oligomeric states (Fig. 2E), similarly to the result obtained with the protein fragment in the *in vitro* situation (Fig. 1D). An altered migration pattern compared to the wild-type was also observed for the Y74D variant with shifts in complex sizes and relative complex abundances, supporting the idea that these amino acids are involved in a crucial assembly interface within the CCDC127 oligomer (Fig. 2E). In a complementary experiment, we performed chemical cross-linking analyses in cell lysates, using the amine-specific crosslinker disuccimidyl-glutarate (DSG), followed by denaturing SDS-PAGE (Fig. 2F). Both full-length CCDC127 and an N-terminal variant containing the CC but lacking the C-terminal domain (CCDC127^1-134^), could be crosslinked to a band representing a dimer.

### The C-terminal helical bundle forms a dimer via a hydrophobic interface

We were not able to obtain crystals of the CCDC127 C-terminal domain, and therefore predicted its architecture with Alphafold3 (AF3) (Abramson *et al*, 2024). In the resulting high-confidence model, the C-terminal region formed an elongated four-helix bundle to which we refer from here on as the ‘C-terminal helical bundle’ (CHB). C174 and C219 form a disulfide bond, linking helices α1 and α4 (Fig. 3A). Notably, the predicted CHB contains two prominent hydrophobic surface areas that were partly conserved on the amino acid level, but completely identical in their biophysical properties (Figs. 3B, C, S4). A peripheral hydrophobic surface contains amino acid residues L137, Y141, L233, L236, Y237, Y240, L243, V244, L247, F250 and I257. At the tip of the predicted CHB, amino acid residues L176, F177, V222 and W223 formed a second extensive hydrophobic surface patch (Fig. 3B), which appeared to be stabilized by the adjacent disulfide bond (Fig. 3A).

**Figure 3:**
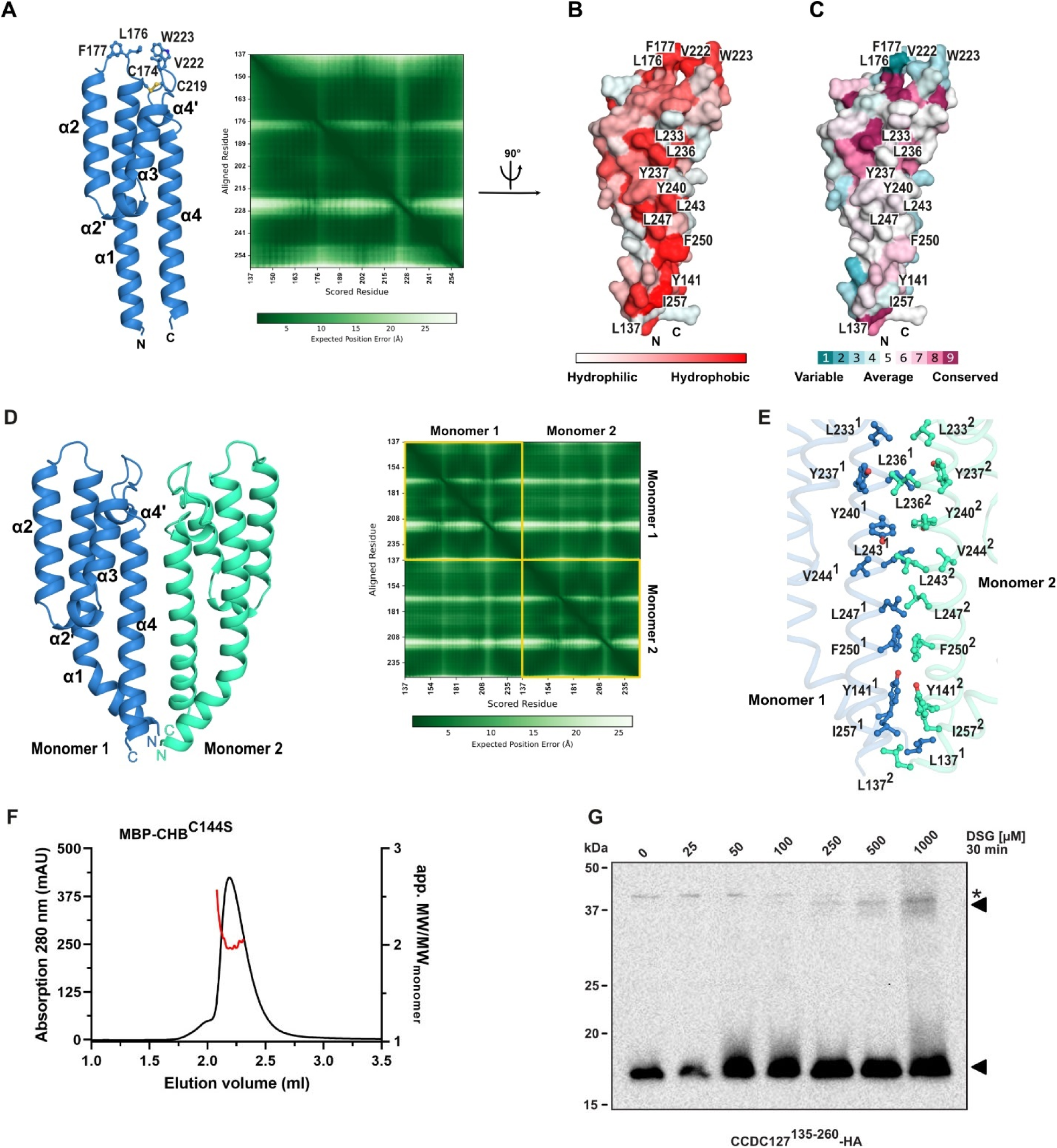
AlphaPulldown identifies CCDC127 C-terminal helical bundle (CHB) as a dimerization domain. **A.** Alphafold3 prediction of the monomeric CHB and the respective predicted aligned error (PAE)-plot. **B, C.** Structural analysis of the surface hydrophobicity (B) and the surface conservation (B) of the monomeric CHB. Residues forming both the hydrophobic patches are labeled. **D.** Structure of the AlphaFold3-predicted dimeric CCDC127 CHB (residue 137-260) and respective PAE-plot. **E.** Dimer interface of the dimeric CHB domain. The interface is characterized by hydrophobic contacts of 11 amino acids (L137, Y141, L223, L236, Y237, Y240, L243, V244, L247, F250, I257). **F.** Analytical SEC-RALS analysis of MBP-tagged C144S (MBP-CHB^C144S^, residue 137-260) variant on a Superose 6 5/150 SEC column. The graphs show the absorption at 280 nm on the left y-axis (black line), the apparent molecular weight divided by the molecular weight of the monomer on the right y-axis (red line) and the elution volume in ml on the x-axis. **G.** CCDC127^135-260^ -HA can be crosslinked in HEK293 cells using the crosslinker DSG with increasing concentrations.

**Figure 4:**
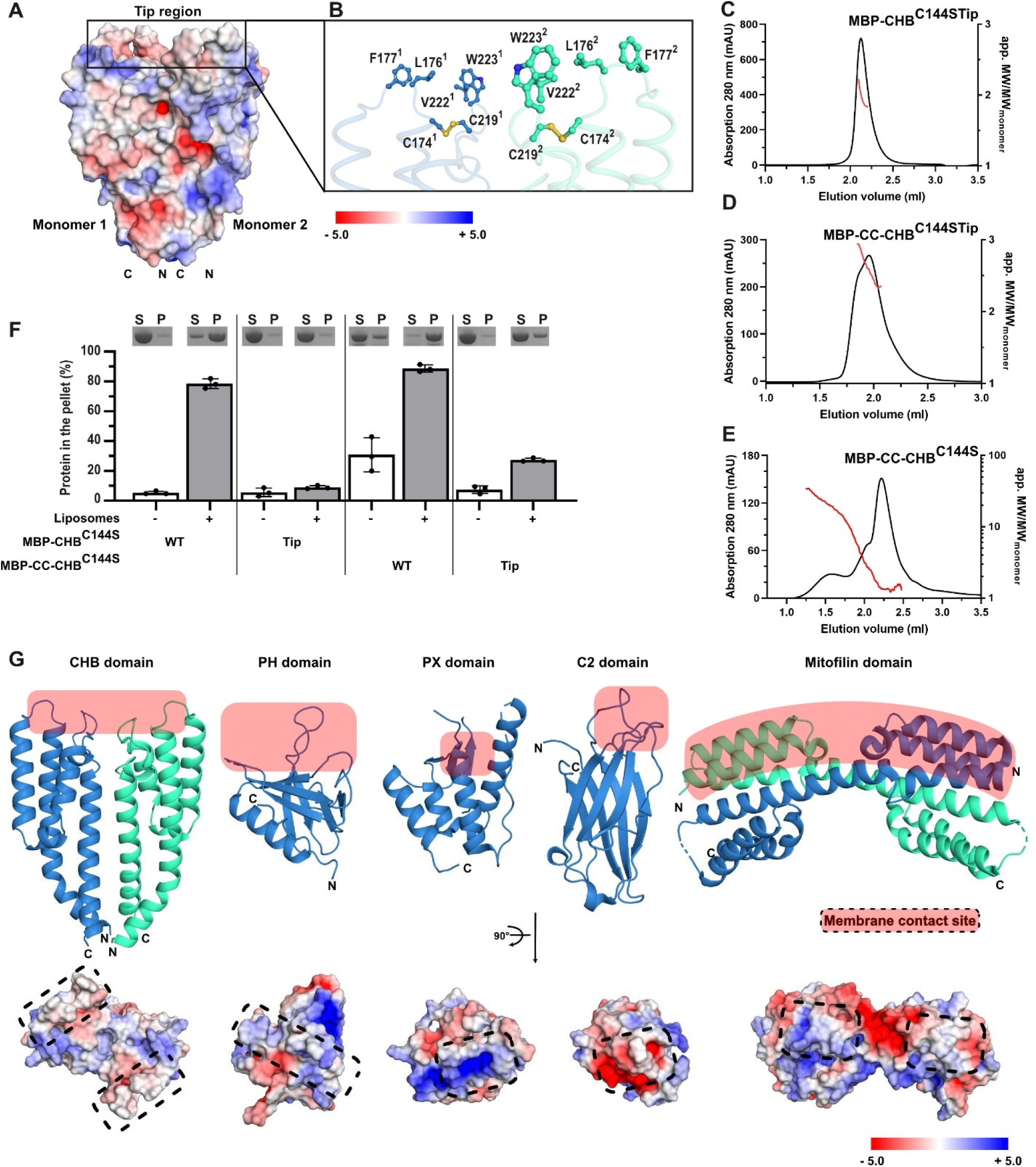
The dimeric CHB is a peripheral membrane-binding domain. **A.** Surface charge distribution of the CHB domain dimer. **B.** Close up of the tip-region of the dimeric C-terminal helical bundle (CHB, residue 137-260). This region contains the important disulfide bond C174-C219 and a hydrophobic patch (L176, F177, V222 and W223). **C.** Analytical SEC-RALS analysis of MBP-tagged C-terminal helical bundle C144S (MBP-CHB^C^, residue 137-260) and a variant containing in addition the L176S/F177S/V222S/W223S (MBP-CC-CHB^C144STip^) alterations, as described in Fig. 3F. **D.** Analytical SEC-RALS analysis for complete IMS domain MBP-CCDC127 C144S containing the L176S/F177S/V222S/W223S quadruple amino acid change (MBP-CC-CHBC144STip). **E.** Analytical SEC-RALS analysis for complete IMS domain MBP-CCDC127 C144S (MBP-CC-CHB^C144S^). **F.** SDS-PAGE analysis and quantification of liposome co-sedimentation assay of MBP-tagged C-terminal helical bundle C144S (MBP-CHB^C144S^) and MBP-tagged complete IMS domain CCDC127 C144S variant without (MBP-CC-CHB^C144S^) and with amino acid exchanges in the tip region (L176S/F177S/V222S/W223S, MBP-CC-CHB^C144STip^) in the absence and presence of liposomes. SN: supernatant; P: pellet. **G.** Structural comparison of the CHB to other peripheral membrane binding domains PH domain of human dynamin (pdb 5A3F, (Reubold *et al*, 2015)), PX domain of human SNX9 (pdb 2RAI, (Pylypenko *et al*, 2007)), the C2 domain of human synaptotagmin (pdb 2R83, (Fuson *et al*, 2007)) and the dimeric mitofilin domain of *Chaetomium thermophilum* Mic60 (pdb 7PV1, (Bock-Bierbaum *et al*., 2022). The membrane contact sites are highlighted by slight red background or dotted lines. The lower panel shows the structural analysis of the surface charge distribution.

The CHB alone was mostly insoluble in expression trials, which may be related to the surface-exposed hydrophobic patches leading to aggregation. However, we succeeded in expressing and purifying a maltose-binding protein (MBP) fusion of a C144S variant of the CHB (MBP-CHB^C144S^, residue 137-260) (Fig. S1E). Interestingly, the fusion protein eluted in two peaks in the final size exclusion chromatography (SEC) step (Fig. S1F). To validate the predicted model, we verified the presence of the disulfide bond between C174 and C219 in our recombinantly expressed protein constructs by mass spectrometry under non-reducing and reducing conditions. All constructs showed a shift in molecular weight of minus 2 Da under non-reducing conditions, indicating proper disulfide bond formation (Fig. S2, S3). Consequently, we purified the constructs in the absence of reducing agents to maintain the integrity of the C174-C219 disulfide bond. Under these oxidizing conditions, the C144S amino acid exchange prevented the formation of covalent oligomers of the CHB, as occasionally observed for the wild-type construct.

We reasoned that the hydrophobic surface patches may serve as an interaction platform for other proteins in the IMS. To identify such potential interaction partners, we employed AlphaPulldown (Yu *et al*, 2023) in an *in silico* screening approach with the CHB as the prey and 612 mitochondrial proteins as baits (Fig. S5A) (Rath *et al*, 2021). Surprisingly, the CHB itself was predicted with high confidence as the most highly ranked interaction candidate (Fig. 3D, Table S3). The predicted CHB homo-dimer featured a two-fold symmetry via the peripheral hydrophobic surface, with an extensive buried surface area of 1350 Å^2^ (Fig. 3E). Since the N-termini of both CHB monomers were pointing next to each other in the same direction, the CHB dimeric prediction is consistent with the experimentally observed parallel coiled-coil dimer, allowing a direct connection of the two dimeric domains.

To analyze the assembly status of the MBP-CHB^C144S^ variant, we used SEC experiments paired with right-angle light scattering (SEC-RALS). The first peak from the initial purification represented various forms of a higher-order oligomer, whereas the second peak corresponded to a dimeric species (Fig. 3F, S1H). These results are consistent with the predicted dimer model of the CHB. Due to its large size and the involvement of eleven residues (Fig. 3E), we refrained from mutating the predicted dimerization interface. Dimer formation of the CHB was also in line with DSG-mediated crosslinking experiments in cell lysates, which led to a weak but reproducible band on denaturing SDS-PAGE corresponding to a dimer (Fig. 3G).

In addition to the CHB, several other proteins were suggested as potential interaction partners of the peripheral hydrophobic surface of CCDC127 in the AlphaPulldown experiment (Fig. S5B, Table S3). Most notably, several members of the mitochondrial fission regulator 1 family (MTFR1, MTFR2, MTFR1L) were predicted to interact with the CHB via an amphipathic helix. However, these predictions should be considered with caution, as MTFR1 was recently shown to reside in the MOM, with the amphipathic helix exposed to the cytosol (Tilokani *et al*, 2022).

### The CHB is a peripheral membrane-binding domain

Dimerization of the CHB brought the two hydrophobic tip regions in close proximity to each other, forming a contiguous surface (Fig. 4A, B). To explore whether this surface may facilitate further assembly of the CHB, we generated a CHB construct (MBP-CHB^C144S^) with the additional L176S, F177S, V222S and W223S quadruple substitution in the tip region (MBP-CHB^C144STip^). Compared to the previously described C144S-only construct (Fig. 3F, G), this construct did not form higher-order oligomers in SEC-RALS analysis, leading to an exclusively dimeric species (“Tip variant”, Fig. 4C). Similarly, an MBP-tagged construct of the complete IMS domain of CCDC127 containing the quadruple tip substitution and the C144S mutation (MBP-CC-CHB^C144STip^, residues 48-260) formed an assembly with 2-3 monomers (Fig. 4D). We also generated the MBP-CC-CHB construct containing the C144S mutation with an intact tip region. Compared to the tip variant, the resulting fusion protein formed dimers and higher-order oligomers, which may indicate a role of the tip region in mediating higher order assembly (Fig. 4E).

The hydrophobic nature of the tip region appeared reminiscent of the membrane-binding sites of peripheral membrane proteins (Tubiana *et al*, 2022). We therefore explored the capacity of the MBP-tagged CHB to interact with membranes, using co-sedimentation experiments with Folch liposomes (comprised of a lipid extract from bovine brain). In the absence of liposomes, the MBP-tagged CHB remained soluble in the supernatant (Fig. 4F, S1G). Strikingly, the C144S variant could be efficiently co-sedimented with Folch liposomes, while sedimentation of the corresponding variant in the tip region was greatly diminished (Fig. 4F). In a similar fashion, liposome-dependent co-sedimentation was also observed for MBP-tagged complete IMS moiety of CCDC127. However, different from the MBP-tagged CHB, some sedimentation was already observed in the absence of liposomes for this construct, in agreement with its observed higher-order assembly formation (Fig. 4F). These experiments indicate a function of the CHB as a novel peripheral membrane-binding protein.

Compared with other peripheral membrane-binding domains, such as PH, PX, C2 or the mitofilin domains (Fig. 4A, G), the central membrane interaction surface of the CHB is more hydrophobic, with a surrounding ring of peripheral polar residues (“polar belt”) potentially interacting with the lipid headgroups. While PH and PX domain typically bind as monomers to membranes, the membrane-binding area of the CHB is extended by dimerization, similar to the mitofilin domain dimer. The dimeric CHB membrane contact surface appears to be flat, whereas the mitofilin domain dimer features a highly curved membrane-binding site.

## DISCUSSION

The first membrane bridging and scaffolding protein machineries have been identified in recent years that protrude and squirm through the mitochondrial IMS, giving rise to the idea of an interconnected network of proteinaceous bundles, coils and fibers that organize the structure and functionality of this rather tiny sub-compartment. Maybe the best characterized component of this putative ‘mitoskeleton’ structure is the MICOS complex with its crista junction shaping and membrane tethering elements. Of note, several auxiliary proteins additionally link MICOS to other mitochondrial functions, like protein import and sorting or mitochondrial DNA (mtDNA) organization and stability. As part of our studies on the MICOS interactome, we report in this work on the identification of a novel MICOS interactor, termed Coiled-Coil Domain-Containing 127 (CCDC127). Our structural analysis of CCDC127 suggests a function as a dynamic membrane bridging contact site protein of the IMS and, thus, as a component of the hypothetical mitoskeleton structure. In support for such a function and in line with results from a companion manuscript (Zarges *et al*, 2025b) and a recent preprint (Hassdenteufel *et al*, 2025), CCDC127 knockout cells displayed altered mitochondrial membrane morphology, with smaller and often irregularly branched cristae (Fig. S6).

Using the available structural data, we propose models of human CCDC127 in the dimeric and oligomeric forms (Fig. 5A, B). The models feature a parallel dimeric assembly of the coiled-coil regions, which are connected by a short linker to the dimeric C-terminal helical bundles. In this configuration, the TM regions can be inserted in parallel into one membrane bilayer, constituting the CCDC127 dimer as the basic structural unit (Fig. 5A). Our biochemical analyses indicate that two CCDC127 dimers may further assemble via a conserved tetramerization interface in the coiled-coil region. When modelling this assembly based on the crystal structure of the isolated coiled-coil domain, the four TMS can insert into the same membrane bilayer, while two dimeric CHBs are located 32 nm away from each other (Fig. 5B). The CHB tip regions may then allow the further assembly of the tetramer into a filamentous structure below the membrane (Fig. 4E). With the currently available data, the appearance and potential function of higher-order CCDC127 assemblies in the mitochondrial IMS are not fully clear and further experimental evidence is required to corroborate the results obtained here.

**Figure 5:**
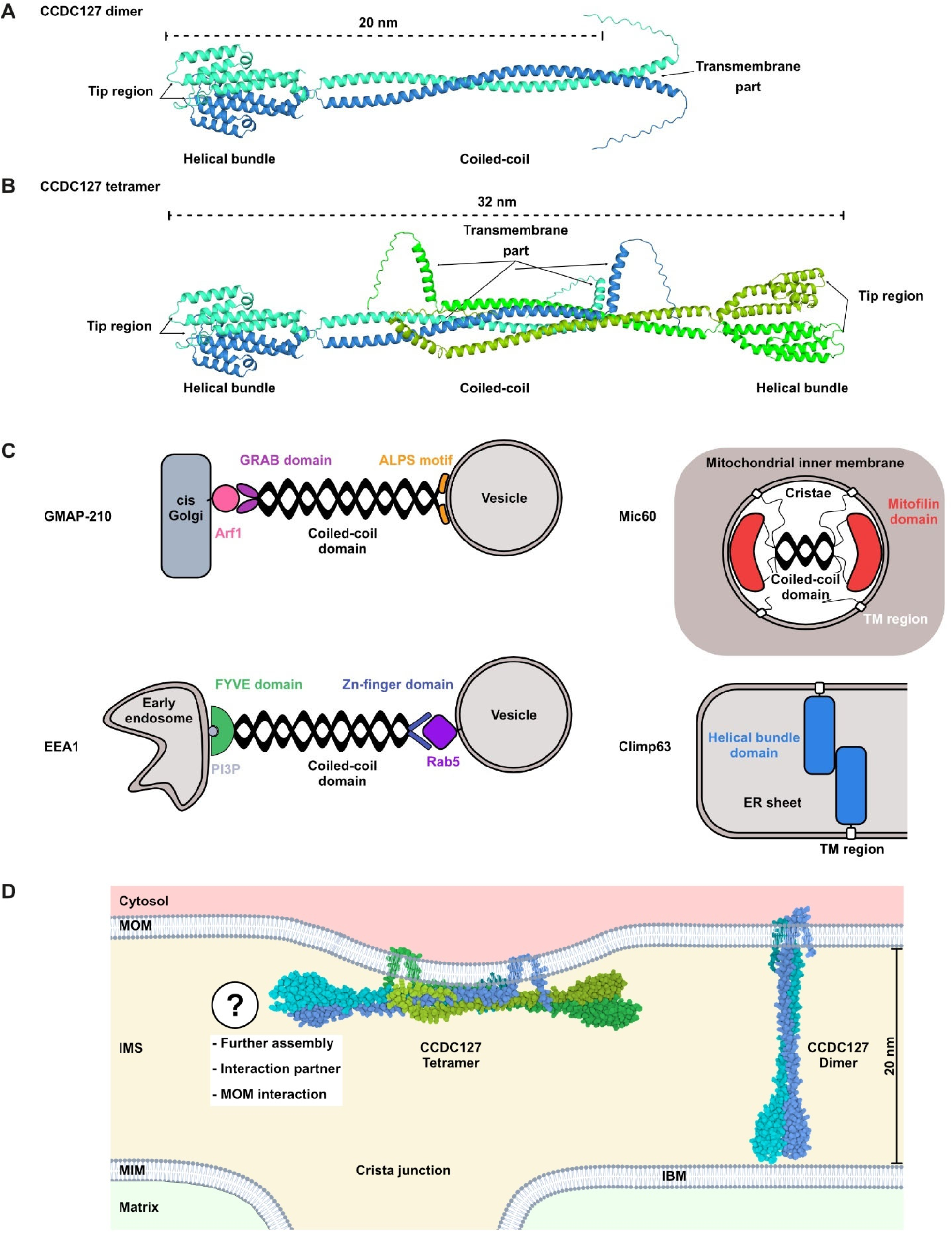
Structural model of CCDC127 as novel membrane contact site protein bridging the IMS. **A.** Alphafold3 prediction of the CCDC127^fl^ dimer. The distance was measured between W44 (representing the last amino acid of the TM part) and W223 at the membrane-interacting tip region. **B.** Model of the CCDC127^fl^ tetramer. This model has been generated using the tetrameric coiled-coil structure derived from x-ray crystallography as template and the Alphafold3 prediction of the full-length dimer. After placing two copies of the dimer, the transmembrane part has been adjusted in order to reach the membranes. The distance was measured between W223 (tip region that interacts with the membrane) from both sides. **C.** Schematic comparison to other membrane contact site proteins: GMAP-210 (Gillingham, 2018), EEA1 (a Golgin protein family member) (Dumas *et al*, 2001; Mishra *et al*, 2010)), Mic60 (Bock-Bierbaum *et al*., 2022) and Climp63 (Xu *et al*., 2023; Zhang & Hu, 2016). **D.** Model for the dimeric and tetrameric arrangement of CCDC127 in the IMS. CCDC127 might span the intermembrane space and connecting both, the MOM and in the MIM. How both the CHB domain dimers interact with membranes in the tetrameric arrange remains an open question and requires further investigation.

In addition, we show that the CHB of CCDC127 comprises a novel peripheral membrane-binding domain. The architecture of the tip region, featuring a hydrophobic core surrounded by positively charged amino acids, closely resembles the membrane-interaction sites of other peripheral membrane-binding modules, such as PH domains (Khurana *et al*, 2023) or the tip region of EHD proteins (Melo *et al*, 2017). While the hydrophobic residues would be inserted into the outer leaflet of the membrane, the polar residues may mediate contacts towards the head groups of polar lipids. In the model of the CCDC127 dimer, the coiled-coil region separates the TMS parts on one side from the C-terminal helical bundles on the other side of the protein, spanning a distance of ∼20 nm (Fig. 5A, D). Notably, the MOM and MIM were shown to be separated by ∼20 nm across large boundary areas (Kühlbrandt, 2015). We therefore envisage that the CCDC127 dimer constitutes a membrane contact site between the MOM and MIM, with the TMS inserted into one membrane and the helical bundles binding to the opposite membrane (Fig. 5D).

We noted that CCDC127 has a closely related architecture compared to other membrane contact site proteins. For example, proteins of the Golgin family also feature parallel dimeric coiled-coil regions and mediate membrane tethering at the Golgi membrane with membrane-binding, TMS or protein interaction motifs at either side of the coiled-coil (Fig. 5C) (Gillingham, 2018; Muschalik & Munro, 2018; Ungermann & Kümmel, 2019). Also, the IMS-exposed protein Mic60 contains an N-terminal TM domain, followed by a tetrameric coiled-coil and a C-terminal mitofilin domain constituting a peripheral membrane-binding site (Bock-Bierbaum *et al*., 2022; Daumke & van der Laan, 2025). In a current model of Mic60 function, the coiled-coil domain traverses crista junctions, whereas the TM domains and mitofilin domains interact with the membrane on both sides of the crista junction (Fig. 5C). In addition, the endoplasmic reticulum-located protein Climp63 (also known as CKAP4) consists of and N-terminal TMS followed by a predicted coiled-coil domain (Xu *et al*, 2023; Zhang & Hu, 2016). Even if the exact architecture of the Climp63 coiled-coil domain remains elusive, initial structural predictions suggest an α-helical bundle built by 4-5 individual helices. Interestingly, Climp63 has been shown to oligomerize via its C-terminal α-helical bundle in a tip-to-tip-like manner, thereby spanning the luminal space in the ER supporting the formation of cisternal structures (Xu *et al*., 2023; Zhang & Hu, 2016). In a similar way, the coiled-coil domains of CCDC127 may act as a spacer between two membranes, with the TMS and the CHB constituting the membrane attachment sites (Fig. 5D).

Together, our biochemical and structural investigation provides the first detailed insights into the molecular architecture of CCDC127. We propose a potential function of CCDC127 as a MICOS-associated membrane tethering unit in the mitochondrial intermembrane space by binding the two opposing membranes at the same time. This idea is in line with a companion manuscript that describes the import mechanism of CCDC127 into mitochondria via the disulfide relay system and identifies functions of CCDC127 in phospholipid homeostasis and mitochondrial shape generation (Zarges *et al*., 2025b). Further corroborating our model, a recent preprint describes the identification of CCDC127 in a screen for mitochondrial proteins impacting on mitochondrial ultrastructure (Hassdenteufel *et al*., 2025). This study provides initial evidence for a role of CCDC127 in cardiolipin metabolism. Further experimental investigation will characterize the detailed role of CCDC127 in maintaining membrane integrity, proper protein distribution or scaffolding of entire membrane patches.

## METHODS

### Cross-linking spatial proteomics

The data shown in Fig. 1A were taken from two previously published XL-MS datasets, both derived from (Zhu *et al*., 2024). Data processing and figure generation followed the same procedures as described in the original publication. Briefly, mitochondria were isolated from HEK293T cells and cross-linked using disuccinimidyl sulfoxide (DSSO) or the enrichable azide-tagged, acid-cleavable disuccinimidyl bis-sulfoxide (DSBSO). DSSO-cross-linked peptides were fractionated by strong cation exchange (SCX). DSBSO-cross-linked peptides were enriched using dibenzocyclooctyne (DBCO)-coupled sepharose beads, and fractionated by size exclusion chromatography (SEC) and high pH (HPH) fractionation. Cross-linked peptides were identified by LC-MS/MS on an Orbitrap Fusion Lumos and analyzed using a stand-alone version of XlinkX. Identifications were filtered at 2% FDR at the unique residue pair level.

### Mammalian cell culture

Human cervical carcinoma (HeLa) cell lines were cultivated in Dulbecco’s Modified Eagle Medium (DMEM) with 10% (v/v) fetal bovine serum (FBS), 1% (v/v) penicillin/ streptomycin (P/S, Gibco), 4.5 g/l glucose and 1% (v/v) L-glutamine (Gibco). Human embryonic kidney cells (HEK293T) were cultivated in DMEM substituted with 4.5 g/l glucose, 4 mM L-glutamine, 1 mM pyruvate, 10% (v/v) fetal bovine serum (FBS), and 50 µg/ml uridine at 37 °C and 5% CO_2_. All cell lines were regularly tested for mycoplasm.

### Transient transfections

Construct of CCDC127-HA expressing C-terminal HA-tagged human CCDC127 under a CMV Promotor (pcDNA3.1(+) backbone) was synthesized by Absea Biotechnology and verified by Sanger Sequencing. Transfections of wild-type HeLa cells with this plasmid were carried out using Lipofectamine (Invitrogen) as per the manufacturer’s instructions. In preparation, cells were seeded 24 h prior to transfection in low confluency. The next day, a mixture of plasmid DNA and Lipofectamine 2000, at a ratio of 1:0.8 (Lipofectamine 2000 volume (μl) to plasmid DNA (μg)), was prepared separately in Opti-MEM. After mixing both solutions briefly, Lipofectamine 2000 was added to the plasmid DNA and the combined mixture was incubated for 20 min at RT before being added dropwise to the cells. After 24 h of transfection, cells were used either for microscopy or other experiments.

### Immunofluorescence

Untransfected and plasmid-transfected HeLa cells were fixed with PBS containing 4% para-formaldehyde (PFA) and 4% sucrose for 10 min at RT. Fixation was quenched by removing PFA and adding PBS containing 0.1 M glycine and 0.1 M ammonium chloride for 10 min at RT. Cells were permeabilized by PBS containing 0.15% Triton X-100 for 10 min at RT and washed with PBS twice. Permeabilized cells were sequentially incubated in blocking buffer (PBS containing 1% bovine serum albumin [BSA] and 6% normal goat serum [NGS]) for 30 min at RT, followed by blocking buffer containing primary antibodies against the HA-tag (rat, Chromotek 7c9-100, 1:200), TOMM20 (mouse, F10, Santa Cruz sc-17764, 1:100) and/or CCDC127 (rabbit, HPA045052, 1:50) for one hour at RT or at 4°C overnight. After three washes in blocking buffer, the cells were incubated in blocking buffer containing highly cross-absorbed secondary antibodies anti-rabbit, CF640R Biotium 20178-1, anti-mouse CF488A Biotium 20014 and anti-rat CF568 Biotium 20092-1 in 1:2000 dilutions for 30 min at RT. After three washes in blocking buffer, the cells were either directly imaged after being stored in PBS or mounted using Pro Long™ Gold (Invitrogen™, P 69) for later imaging.

### Confocal immunofluorescence microscopy

Confocal images were acquired with a Nikon spinning disc microscope (Yokogawa spinning disk CSU-X1). The microscope is equipped with the following lasers and emission filters (Exc. 488 nm (Em. 525/50 nm), Exc. 561 nm (Em. 600/50 nm) and Exc. 638 nm (Em. 700/75 nm)), a 60x oil objective (Plan-Apo, NA 1.40 Nikon), an Andor camera (AU888, 13 μm/pixel) and NIS Elements software. Cells of fixed images were acquired with a final pixel size of 110 nm for the 60x objective and stored using NIS Elements software.

### Generation of cell lines

For the generation of CCDC127 knockout cells, guide RNA sequences targeting the *CCDC127* gene were cloned into the pSpCas9(BB)-2A-GFP (PX458) vector kindly provided by Mike Ryan (Monash University Melbourne, Australia; Addgene plasmid # 48138) and/or Feng Zhang. The guide RNA sequences used were Guide 1: 5’-CACCGTGAGATCATGGCGTGGTACT-3’ and Guide 2: 5’-CACCGACGCCGATTTTCTGAGATCATGG-3’. HEK293T (guide 1) and HEK Flp-IN-T-Rex-293 (guide 2) cells were transfected with the plasmids using polyethylenimine (PEI; Thermo Fisher Scientific). After 24 h, GFP-positive cells were collected via FACS and single-cell clones were seeded into 96-well plates (1 cell per well). Clonal cell lines were screened for the loss of CCDC127 by western blotting.

Cells expressing CCDC127 variants were generated via two different methods. For retroviral transduction using the pBABE vector system, the sequences encoding CCDC127-FLAG, FLAG-CCDC127, CCDC127-K73A or -K74D were cloned into the pBABE-puro vector and co-transfected with pGagPol and pVSVG vectors in high-virus-titre-producing HEK293T helper cells using Lipofectamine LTX, according to the manufacturer’s instructions. HEK293T ΔCCDC127 cells were infected with virus-containing supernatant, selected with puromycin and expression of CCDC127 variants was verified by Western blot analysis. For the complementation of CRISPR clones with either CCDC127-HA, CCDC127 coiled-coil (amino acid 1-134) or C-terminal helical bundle domain (amino acids 135-260), the inducible Flp-In T-REx System was used. These CCDC127 constructs were cloned into the pcDNA5 FRT-TO vector and co-transfected with the pOG44 Vector into the different CRISPR clones by using the transfection reagent FuGene, according to the manufacturer’s guideline. Positive clones were selected with glucose-containing medium (DMEM supplemented with 1 mM sodium pyruvate, 1 x nonessential amino acids, 10% FCS and 500 mg/ml Pen/Strep, 50 µg/ml Uridine) containing 10 µg/ml Blasticidin and 100 µg/ml Hygromycin.

### Cell fractionation

Subcellular localization of CCDC127 was analyzed by cell fractionation. Cells were harvested and washed in PBS before being resuspended in buffer A (83 mM sucrose, 10 mM HEPES/KOH pH 7.2) and disrupted with 10 strokes in a glass homogenizer. After centrifugation at 1,000 x *g* for 5 min at 4 °C, the supernatant and pellet were separated. The pellet (nuclei fraction) was treated with DNase and dissolved in Laemmli buffer. A portion of the supernatant was mixed directly with Laemmli buffer (lysate fraction = total without nuclei). The remaining supernatant containing organelles and cytosol, was centrifuged at 12,000 x *g* for 15 min at 4 °C, and the pellet and supernatant were separated. The pellet was washed in buffer B (320 mM sucrose, 20 mM HEPES/KOH pH 7.2, 1 mM EDTA) and resuspended in Laemmli buffer, whereas the supernatant (cytosol and light membranes) was TCA-precipitated prior to dissolving the pellet in Laemmli buffer. Samples were analyzed by SDS-PAGE and Western blot with antibodies against CCDC127 (Invitrogen PAS-60912), TOMM22 (abcam ab179826), α-Tubulin (abcam ab7291) and Histon H3 (Cell Signaling Technology #9715).

### Alkaline extraction of proteins

To assess the membrane association of proteins, mitochondrial pellets were resuspended in freshly prepared 0.1 M Na_2_CO_3_ pH 11 or pH 11.7 and incubated on ice for 30 min. Half of the sample was kept as the total fraction, whereas the other half was subjected to ultracentrifugation in a TLA45 rotor at 125,000 x *g* for 30 min at 4 °C. The supernatant and pellet fractions were separated. The total and supernatant fractions were TCA precipitated, and all pellets were dissolved in Laemmli-buffer while shaking at 60 °C for 10 min. Samples were analyzed by SDS-PAGE and Western blot with antibodies against CCDC127 (Invitrogen PA5-60912), TOMM22 (abcam ab179826) and HSPA9 (abcam ab227215).

### Proteinase K accessibility assay

Submitochondrial localization and membrane topology of CCDC127 were analyzed by mitochondrial swelling and proteinase K treatment. HEK293T cells expressing either endogenous CCDC127 or N- or C-terminally FLAG-tagged variants were harvested and resuspended in buffer A (20 mM HEPES/KOH pH 7.6, 220 mM mannitol, 70 mM sucrose, 1 mM EDTA, 2 mg/ml BSA and 0.5 mM PMSF) before being homogenized with 20 strokes in a glass homogenizer. After centrifugation at 800 x *g* for 5 min at 4 °C, the supernatant was centrifuged at 15,000 x *g* for 10 min at 4 °C. The resulting pellet containing crude mitochondria, was resuspended in buffer B (20 mM HEPES/KOH pH 7.6, 220 mM mannitol, 70 mM sucrose, 1 mM EDTA and 20 mg/ml BSA). Following a second centrifugation at 15,000 x *g* for 10 min at 4 °C, the mitochondrial pellet was resuspended in buffer B. Protein concentration was determined using the BCA-reagent ROTI^®^Quant and 100 µg of mitochondria were pelleted at 15,000 x *g* for 10 min at 4 °C for each of the four experimental conditions. The pellets were either resuspended in SM buffer (10 mM MOPS/KOH pH 7.2, 320 mM sucrose) or MSM buffer (10 mM MOPS/KOH pH 7.2, 16 mM Sucrose) and incubated for 30 min at 4 °C. From each buffer condition, one sample was treated with 5 µg proteinase K or mock treated with SM buffer for 30 min on ice. Reactions were stopped by the addition of PMSF to a final concentration of 4 mM. All samples were centrifuged at 15,000 x *g* for 5 min at 4 °C, and pellets were washed with SEM buffer (10 mM MOPS/KOH pH 7.2, 320 mM sucrose, 1 mM EDTA) followed by a second centrifugation under the same conditions. Supernatants were discarded and pellets resuspended in 100 µl Laemmli buffer, incubated for 10 min at 65 °C and analyzed by SDS-PAGE and Western blotting with antibodies against CCDC127 (Invitrogen PA5-60912), FLAG (Sigma F7425), TOMM22 (abcam ab179826), MIC10 (homemade serum raised in the Pfanner/van der Laan labs #5032) and HSPA9 (abcam ab227215).

### Crosslinking of CCDC127 with disuccinimidyl glutarate (DSG)

In order to check whether CCDC127 can be crosslinked in intact cells, HEK cells were seeded in a 12-well plate until they reached 90% confluency. Cells were placed on ice and washed twice with ice-cold PBS. Afterwards the DSG-PBS solution with different concentrations of DSG (0, 25, 50, 100, 250, 500, 1000 µM) was added carefully. Incubation occurred for 30 min. The reaction was stopped by removing the crosslinker and washing the cells with 20 mM Tris pH 7.5, and incubation in 20 mM Tris pH 7.5 for 15 min at RT. Cells were scratched off and sedimented at 500 x *g* for 5 min at 4 °C and resuspended in 20 µl 1x reducing SDS sample buffer. The samples were boiled for 10 min at 96 °C and analyzed by Western Blot.

### Transmission electron microscopy

For transmission electron microscopy analysis of ultrathin sections approximately 10,000 HEK293T cells were seeded onto poly-l-lysine-coated 1.4 mm sapphire discs (Leica) and incubated for 24 h at 37 °C and 5% CO_2_. Subsequently, the cells were vitrified in a high-pressure freezing system (EM PACT2; Leica) and embedded in Lowcryl (Polysciences). All of the samples were processed in an automatic freeze-substitution apparatus (AFS2; Leica) and transferred into the precooled (−130 °C) freeze-substitution chamber of the AFS2. The temperature was increased from −130 to −90 °C over a 2 h period. Cryo-substitution was performed in anhydrous acetone and 2% water. The temperature was increased linearly from −90 °C to −70 °C over 20 h, and from −70 °C to −60 °C over 20 h. Increasing concentrations of Lowicryl (50%, 75% and 100%) were added stepwise to the samples at 1 h intervals followed by a 5 h incubation in 100% Lowicryl. Polymerization was carried out under ultraviolet light for 24 h followed by a slow increase of temperature over 15 h to 20 °C. Ultrathin (70 nm) sections were cut using an ultramicrotome (EM UC7; Leica), collected on Pioloform-coated copper grids, stained with uranyl acetate and lead citrate, and analysed with a Tecnai G2 Biotwin electron microscope (Thermo Fisher Scientific). The TEM images were acquired using the Olympus iTEM 5.0 image software (build 1243).

### Recombinant expression and purification of recombinant proteins

Constructs of human CCDC127 (CCDC127; UniProt ID: Q96BQ5) coiled-coil domain (CCDC127^CC^, residues 48-120) were cloned into the tailor-made vector pSKB, leading to an N-terminal His_6_-tagged and a human rhinovirus (HRV)-3C protease cleavable fusion construct. Constructs of the C-terminal helical bundle (CHB, residues 137-260) and the complete IMS domain of CCDC127 (CC-CHB, residues 48-260) have been cloned into the tailor-made vector pCrystMas, resulting in an N-terminal maltose binding protein (MBP) fusion construct. All variants of CCDC127 (CCDC127^CC^ D69A, L70D, K73A, Y74D, MBP-CHB and MBP-CC-CHB C144S and C144S/L176S/F177S/V222S/W223S) were generated using site directed mutagenesis (Liu & Naismith, 2008).

The expression plasmid for CCDC127^CC^ was freshly transformed into *E. coli* Rosetta R2 (DE3) cells and protein expression was carried out in terrific broth containing 50 µg/ml kanamycin and 34 µg/ml chloramphenicol. The cultures were grown at 37 °C and 80 rpm until OD_600_ reached 0.7, protein expression was subsequently induced by the addition of 300 µM isopropyl β-D-1-thiogalactopyranoside (IPTG) and incubated at 20 °C for another 16 h. The next day, the cells were centrifuged at 4,000 x *g*, collected and frozen at -20 °C.

The expression plasmids for the MBP-CHB and MBP-CC-CHB variants were freshly transformed into *E. coli* SHuffle® T7 Express cells (DE3; New England Biolabs) and further treated as mentioned in the manufacturer’s guidelines. Protein expression, however, was induced with 300 µM IPTG.

Cells expressing CCDC127^CC^ were diluted in lysis buffer (50 mM HEPES/NaOH pH 7.5; 500 mM NaCl, 20 mM imidazole, and 1 mg/ml DNase I (Roche)) before lysis using a microfluidizer (Microfluidics). To remove insoluble parts, the solution was centrifuged at 100,000 x *g*, 4 °C and 45 min. The cleared supernatant was further loaded onto a prepacked Ni^2+^ -Sepharose High Performance IMAC resin (Cytivia)-containing gravity flow column charged with 100 mM nickel sulphate and equilibrated in lysis buffer. The column was extensively washed using lysis buffer and bound proteins eluted with lysis buffer containing 50-and 500 mM imidazole, respectively. To remove the N-terminal His_6_-tag, the protein was treated with recombinant and His_6_-tagged HRV-3C-protease during overnight dialysis at 4 °C against dialysis buffer (20 mM HEPES/NaOH pH 7.5; 500 mM NaCl and 25 mM imidazole). A second Ni^2+^ - sepharose-containing column was used to separate the cleaved from the non-cleaved protein and the protease. Finally, a size-exclusion chromatography (SEC) using a S200 column (Cytivia) and SEC buffer (20 mM HEPES/NaOH pH 7.5 and 500 mM NaCl) was applied to separate pure protein from aggregates.

The MBP-CHB and MBP-CC-CHB variant expressing cells were diluted in MBP lysis buffer (50 mM HEPES/NaOH pH 7.5; 500 mM NaCl, 1 mg DNase I (Roche) and protease inhibitor 4-(2-aminoethyl) benzenesulfonyl fluoride hydrochloride (AEBSF, AppliChem) and lysed using sonication (Bändelin Sonoplus). After centrifugation, the cleared supernatant was further loaded onto a prepacked Dextrin Sepharose High Performance resin (Cytivia)-containing gravity flow column equilibrated in MBP lysis buffer. The column was extensively washed using MBP lysis buffer, and bound proteins were eluted with MBP lysis buffer containing 10 mM maltose. The eluted protein was applied onto a SEC using a Superose 6 column (Cytivia) equilibrated with buffer SEC^MBP^ (20 mM HEPES/NaOH pH 7.5, 300 mM NaCl). Pure proteins were flash frozen in liquid nitrogen and stored at -70 °C. The respective variants have been treated in the exact same way as described above.

### Crystallization, data collection, refinement and other tools

Diffraction quality protein crystals of CCDC127^CC^ grew from an initial crystallization condition containing 0.2 M (NH_4_)_2_SO_4_, 25% PEG 4000, 0.1 M sodium acetate pH 4.6 and have been set up with the vapor diffusion method in 96-well sitting drop format at 20 °C using an automated dispensing robot (Art Robbins Instruments). Crystals have been fished out of the drop and flash-cooled in liquid nitrogen prior to data collection.

Diffraction data were collected at -173 °C and 0.918 Å on beamline BL14.1 operated by the Helmholtz–Zentrum Berlin at the BESSY II electron storage ring (Berlin–Adlershof, Germany) (Mueller *et al*, 2025) and indexed, integrated and scaled with XDSAPP (Sparta *et al*, 2016). The structure of CCDC127^CC^ was solved by molecular replacement using Phaser-MR from the PHENIX suite (McCoy *et al*, 2007; Terwilliger *et al*, 2008) and a truncated and poly-alanine stubbed version of a hsCCDC127 (Uniprot accession code: Q96BQ5) model derived from AlphaFold2 (Jumper *et al*, 2021; Varadi *et al*, 2022). AutoBuild was further used to obtain the initial model. The protein crystallized in space group P3_2_21 (154) with six monomers in the asymmetric unit and appeared to have pseudo-translational defects (tNCS), leading to significantly higher R-values of the final model. Refinement was carried out using iterative steps of manual model building in Coot (Emsley *et al*, 2010) and maximum likelihood refinement with individual B-factors, TLS and secondary structure restraints using phenix.refine (Afonine *et al*, 2012). Final structure validation was carried out with MolProbity (Williams *et al*, 2018) and wwPDB Validation Service (https://validate.wwpdb.org). All statistics for data collection and refinement, as well as the corresponding PDB code, can be found in Supplementary Table1 and 2.

The dimer interface of the C-terminal domain (CHB) has been calculated using PDBePISA (Krissinel & Henrick, 2007). The surface conservation plot was created using the ConSurf Server (Ashkenazy *et al*, 2016) with standard settings and multiple sequence alignments using Clustal Omega (Madeira *et al*, 2024). Figures were prepared with PyMOL (The PyMOL Molecular Graphics System, Version 2.5.5 Schrödinger, LLC).

### Analytical ultracentrifugation

The crystalized construct CCDC127^CC^, as well as the indicated mutants, were analyzed at protein concentrations of 1 mg/ml in 20 mM HEPES/NaOH pH 7.5, 500 mM NaCl using a Beckman Optima XL I analytical ultracentrifuge equipped with an An60Ti rotor and double sector cells. Sedimentation velocity measurements were carried out overnight at 20 °C and at a rotor speed of 40,000 rpm. The absorbance data were recorded at a wavelength of 280 nm and in time intervals of 5 min. Sedimentation coefficient distributions *c(s)* were determined with the program Sedfit (Schuck, 2000). The protein partial-specific volume, buffer viscosity and buffer density were calculated using the software Sednterp (Laue *et al*, 1992) . Figures were created with GUSSI (Brautigam, 2015).

### Analytical size exclusion chromatography (SEC)

Analytical SEC was performed using 100 μl of a protein solution (3 mg/ml) on a Superdex S200 10/300 (Cytivia) column at a flow rate of 0.5 ml/min at 4°C. The running buffer contained 50 mM HEPES/NaOH pH 7.5, and 500 mM NaCl.

### Analytical size exclusion chromatography coupled to right-angle light scattering (SEC-RALS)

Analytical SEC-RALS was performed using a Superose 6 5/150 column (Cytivia) and a RALS instrument (Malvern) equilibrated overnight against SEC buffer (20 mM HEPES/NaOH pH 7.5, 150 mM NaCl). The run was performed with 80 μl of a 1 mg/ml protein solution at a flow rate of 0.2 ml/min at 20 °C. The analysis was done by using the Agilent Offline and Malvern Omnisec software.

### Isolation of mitochondria from human cell lines

HEK293T cells were harvested from three to five 15 cm dishes and washed with PBS. The cell pellets were resuspended in buffer A (83 mM sucrose, 10 mM HEPES/KOH pH 7.2) and homogenized in a glass homogenizer. An equal volume of buffer B (250 mM sucrose, 30 mM HEPES/KOH pH 7.2) was added, and the homogenate was centrifuged at 1000 x *g* for 5 min at 4°C. The supernatant was subsequently centrifuged at 12,000 x *g* for 10 min at 4 °C. The resulting pellet was resuspended in buffer C (320 mM sucrose, 10 mM Tris-HCl pH 7.4), and the BCA-reagent ROTI^®^Quant was used to measure the protein concentration.

### Analysis of mitochondrial protein complexes by BN-PAGE

Mitochondrial protein complexes were analyzed by blue native-PAGE (BN-PAGE). Isolated mitochondria were solubilized for 30 min on ice in solubilization buffer (1% [w/v] digitonin, 20 mM Tris-HCl pH 7.4, 0.1 mM EDTA, 50 mM NaCl, 10% [v/v] glycerol, and 1 mM PMSF). After a clarifying spin at 15,000 x *g* for 10 min at 4 °C, loading dye (5 % Coomassie blue G, 500 mM ε-amino n-caproic acid in 100 mM Bis-Tris pH 7.0) was added to the supernatant, and samples were loaded onto a 4-13% BN-PAGE. Protein complexes were analyzed by Western blot with an antibody against CCDC127 (PA5-60912).

### Top-down mass spectrometry

Protein intact mass analyses were conducted on an Agilent 1290 Infinity II UHPLC system coupled to an Agilent 6230B time-of-flight (TOF) LC/MS instrument equipped with an AJS (Agilent Jet Stream Technology) ion source operated in positive ion mode. Protein samples were desalted using a Zorbax 300SB-C3 guard column (2.1 × 12.5 mm, 5 μm). Approximately 0.6 μg of sample was injected for each analysis. LC/MS parameters were adapted from Chalk *et al*. (Chalk, 2017). The ion source was operated with the capillary voltage at 4000 V, nebulizer pressure at 50 psi, drying and sheath gas at 350 °C, and drying and sheath gas flow rate at 12 and 11 L/min, respectively. The instrument ion optic voltages were as follows: fragmentor 290 V, skimmer 65 V, and octupole RF 750 V. MS data were analyzed using the Protein Deconvolution feature of the MassHunter BioConfirm Version 10.0 software (Agilent) that uses the Maximum Entropy algorithm for accurate molecular mass calculation. Deconvolution was performed between mass range of 800 to 2,500 m/z, using peaks with a ratio of signal to noise greater than 30:1. The deconvoluted mass range was set at 25 to 75 kDa and the step mass was 1 Da.

### Liposome co-sedimentation assay

Folch lipids (total bovine brain lipids fraction I, Sigma-Aldrich) were dried under an argon stream, dissolved in 20 mM HEPES/NaOH (pH 7.5), 150 mM NaCl and incubated for 1 h at RT. 40 µl of a reaction mixture containing liposomes (0.6 mg/ml) and 5 μM protein was incubated for 30 min at RT and centrifuged at 200,000 x *g* for 16 min at 20°C. The respective supernatant and pellet fractions were analyzed by SDS-PAGE, and the protein bands were quantified using ImageJ (version 1.50i) (Schneider *et al*, 2012).

### In silico protein-protein interaction search

AlphaPulldown (Yu *et al*., 2023) was used to screen for potential hsCCDC127 interaction partners *in silico*. In this, the C-terminal helical domain of hsCCDC127 (CCDC127^137-260^) as monomer was used as prey and all mitochondrial proteins, that either locate in the MIM, MOM or IMS (MitoCarta3.0) (Rath *et al*., 2021), were used as potential interaction partners (612 proteins). For the interaction screen, the pulldown mode was used and a final cutoff for the inter-chain PAE of 5 was applied as initial filter criteria. The remaining complexes (95) were cross-validated using Alphafold3 structure prediction (Abramson *et al*., 2024) and finally ranked according to their iptm and ptm + iptm scores.

### Generation of CCDC127 full-length models

The dimer of full-length CCDC127 was predicted using Alphafold3 (AF3) without any further corrections. To fully assemble the CCDC127 tetramer, the coiled-coil domains of two AF3-predicted dimers have been superimposed onto the crystal structure of the central coiled-coil domain and adjusted to avoid clashes. In order to orient the N-terminal transmembrane regions in this oligomeric assembly, a kink was introduced between residue Ile43 and Arg46.

## ACKNOWELEDGEMENT

We thank Yvette Roske for help with crystallographic data collection, the entire MX group at BESSYII for support during data collection at beamlines MX14.1, MX14.2, or MX14.3 and Anja Schütz at the MDC Protein Production & Characterization Technology Platform (https://www.mdc-berlin.de/protein-production-characterization) for providing training and technical support during mass spectrometry analyses. We thank the Scientific Computing technology platform at the MDC for providing access to the Max-Cluster.

## AUTHOR CONTRIBUTION

T.B.B. designed CCDC127 constructs, grew crystals, solved protein structures and performed biochemical experiments with support from C.B.. K.C.A. and M.L. performed CCDC127 localization experiments. K.v.d.M., A.A., K.N., and S.J. performed CCDC127 localization studies and structure-based functional experiments. A.K.N. together with TBB set up and analyzed the AlphaPulldown runs. Y.Z. performed cross-linking MS experiments on isolated mitochondria and provided the figure. N.C. performed analytical ultracentrifugation experiments. C.Z. and J.R. performed cross-linking of CCDC127 in intact cells. T.B.B., K.v.d.M, M.v.d.L., and O.D. designed research and interpreted structural and biochemical data. T.B.B., K.v.d.M., M.v.d.L. and O.D. wrote the manuscript with inputs from all authors.

## DATA AVAILABILITY

All data are available in the main text or the supplementary materials. The atomic coordinates of CCDC127^48-120^ were deposited in the Protein Data Bank with accession numbers 9REU.

---

**Figure S1:**
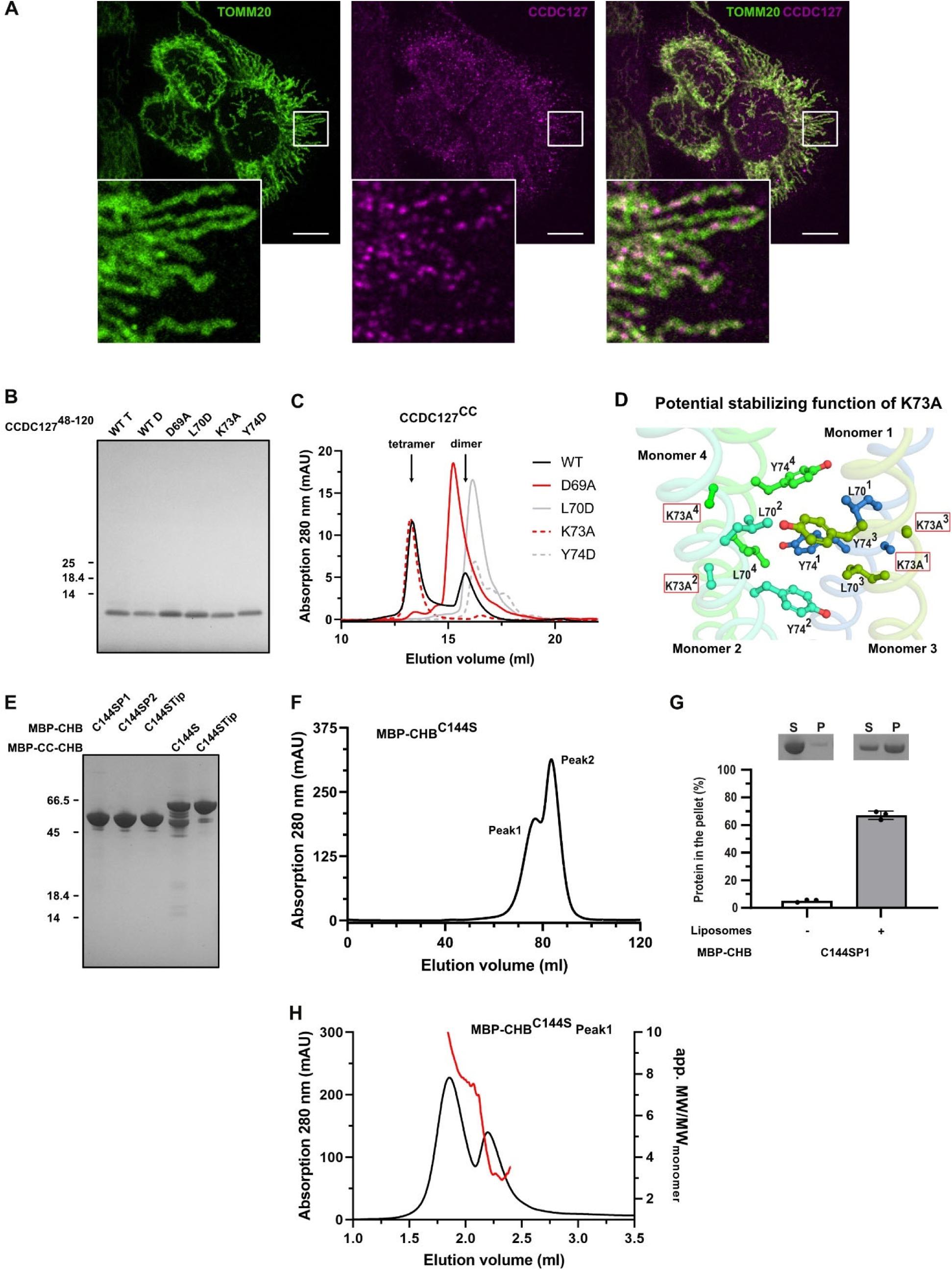
Biochemical and structural analysis of human CCDC127 constructs. **A:** Confocal dual-color microscopy of TOMM20 (left, green), endogenous CCDC127 (middle, magenta) and merged (right) in PFA-fixed un-transfected Hela cells. Insert and scale bar: 10 μm. **B:** Purity of WT coiled-coil domain (CCDC127^CC^, residue 48-120) and variants (D69A, L70D, K73A and Y74D). T – tetramer; D – Dimer in gelfiltration (see Fig. 1D). **C.** SEC running on S200 10/300 SEC column of wildtype and different hsCCDC127 coiled-coil domain (CCDC127^CC^) variants (D69A, L70D, K73A, Y74D). Shown is the absorption at 280 nm in mAU against the elution volume in ml. The wildtype protein forms dimers and tetramers, whereas the variants exist in smaller oligomeric species, as confirmed by analytical ultracentrifugation analysis (**see** Fig. 2D). **D.** Potential role of K73A in stabilizing the coiled-coil tetramer. The newly introduced methyl group in the K73A may reach the hydrophobic core of the coiled coil leading to further stabilization of the tetramer interface. **E.** SDS-PAGE showing the final purity of recombinant C144S and Tip variant (L176S/F177S/V222S/W223S) MBP-tagged C-terminal helical bundle (MBP-CHB^C144S^, MBP-CHB^C144STip^) and complete IMS domain-containing hsCCDC127 (MBP-CC-CHB^C144S^, MBP-CC-CHB^C144STip^). P1 (Peak 1 from SEC run) and P2 (Peak 2 from SEC run). **F.** SEC on Superose 6 16/600 SEC column (Cytivia) of MBP-tagged C-terminal helical bundle (MBP-CHB^C144S^, residues 137-260). Shown is the absorption at 280 nm in mAU against the elution volume in ml. The protein elutes as two peaks (Peak1 and Peak2). **G.** SDS-PAGE analysis and quantification of liposome co-sedimentation assay of peak 1 from SEC of C144S variant MBP-tagged CHB (MBP-CHB^C144S^) in the absence and presence of liposomes. SN: supernatant; P: pellet. **H.** Analytical SEC-RALS analysis of peaks 1 of MBP-tagged C144S C-terminal helical bundle (MBP-CHB^C144S^, residue 137-260) on a Superose 6 5/150 SEC column. The graphs show the absorption at 280 nm on the left y-axis (black line), the apparent molecular weight divided by the molecular weight of the monomer on the right y-axis (red line) and the elution volume in ml on the x-axis.

**Fig. S2:**
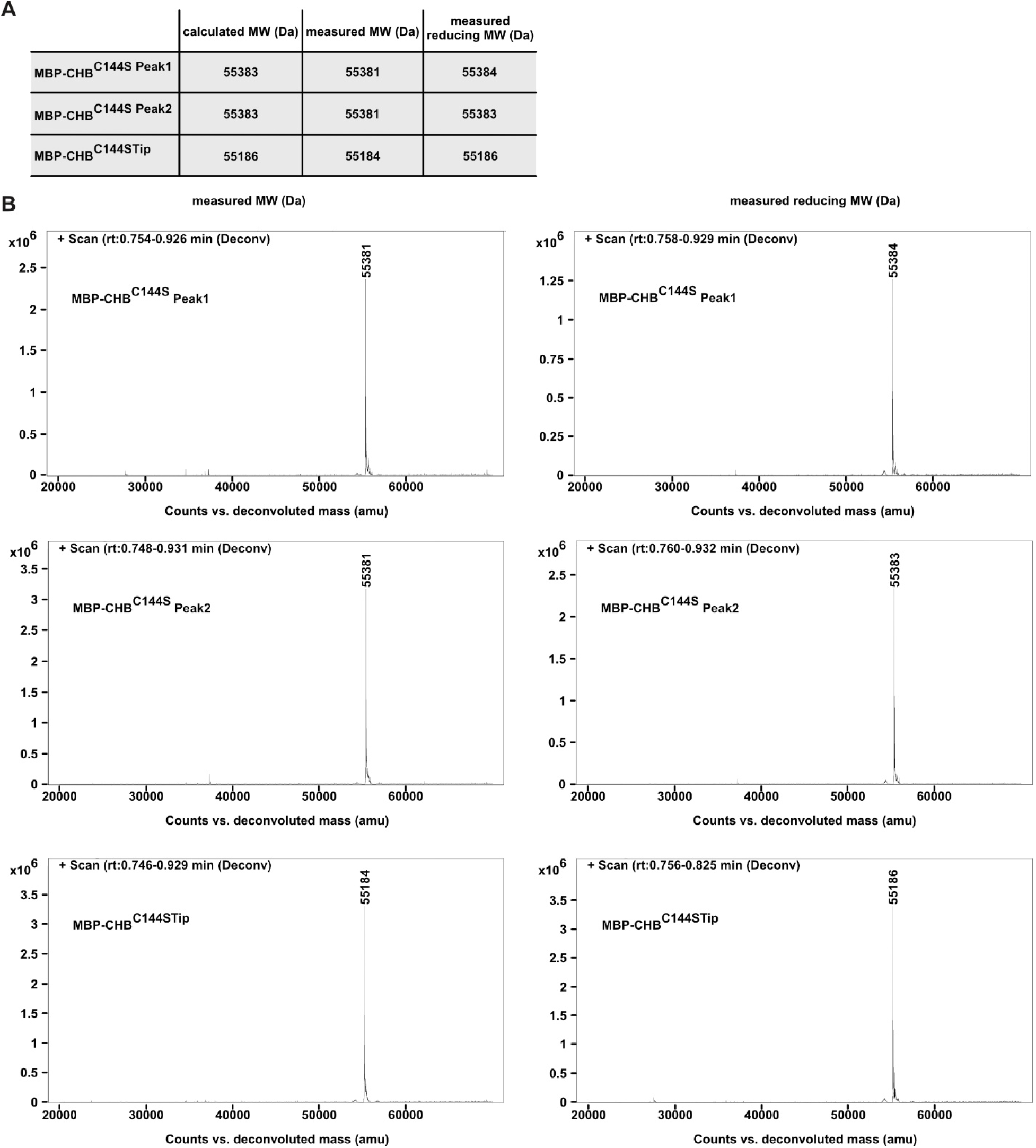
Mass spectrometry analysis of human CCDC127 helical bundle domain constructs. **A and B.** Top-down mass spectrometry measurements under denatured and non-reducing and reducing conditions. Shown are the calculated and the measured molecular mass under denaturing and non-reducing (measured) and reducing (measured reducing) conditions. The shift of - 2 Da is typical for disulfide bond formation (**A**). Deconvoluted spectra collected under non-reducing and reducing (**B**) conditions. Peak 1 and Peak 2 from SEC run (see Figure S1F, H).

**Figure S3:**
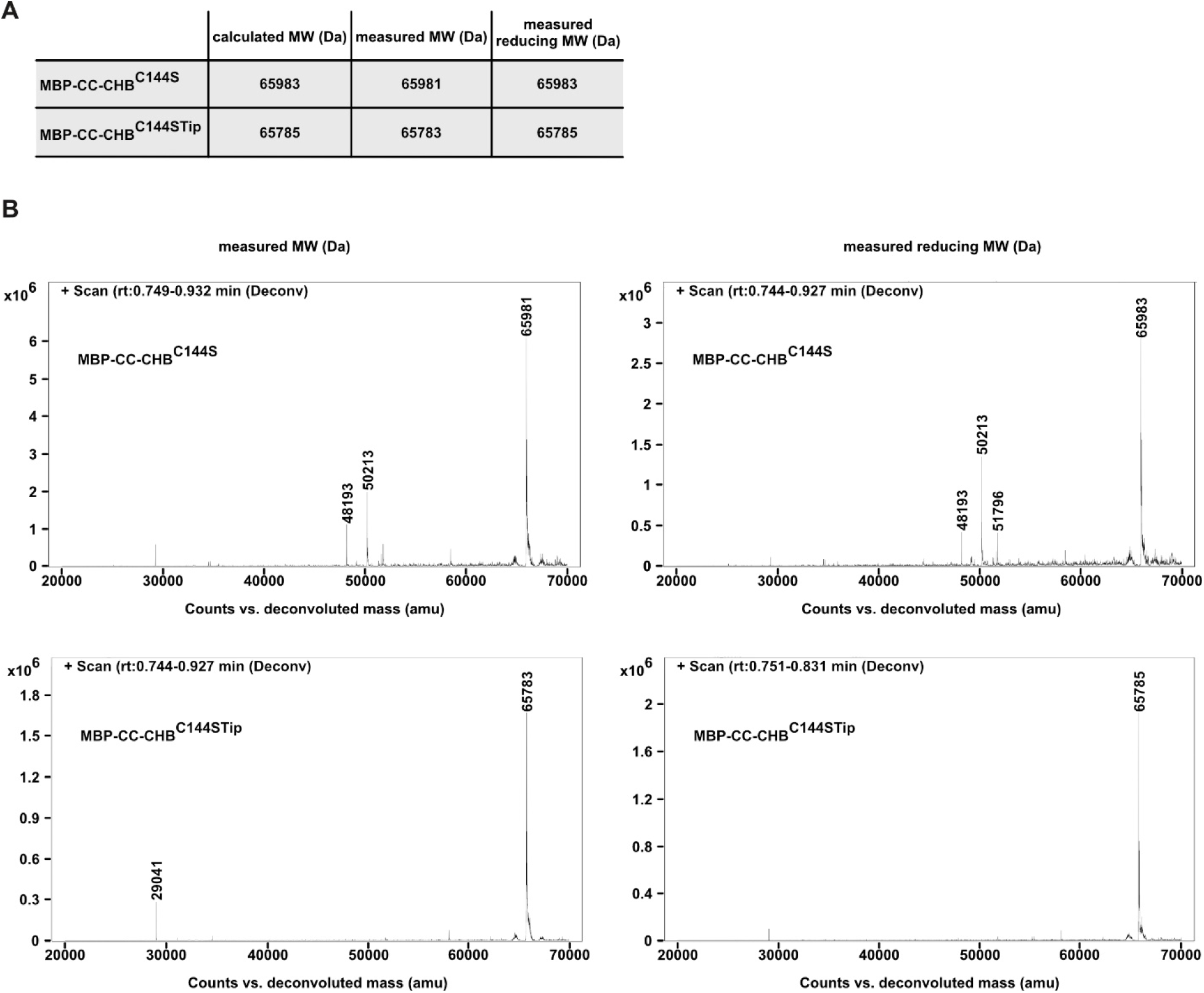
Mass spectrometry analysis of human CCDC127 complete IMS domain constructs. **A and B.** Top-down mass spectrometry measurements under denatured and non-reducing and reducing conditions. Shown are the calculated and the measured molecular mass under denaturing and non-reducing (measured) and reducing (measured reducing) conditions. The shift of - 2 Da represents disulfide bond formation (**A**). Deconvoluted spectra collected under non-reducing and reducing (**B**) conditions.

**Figure S4:**
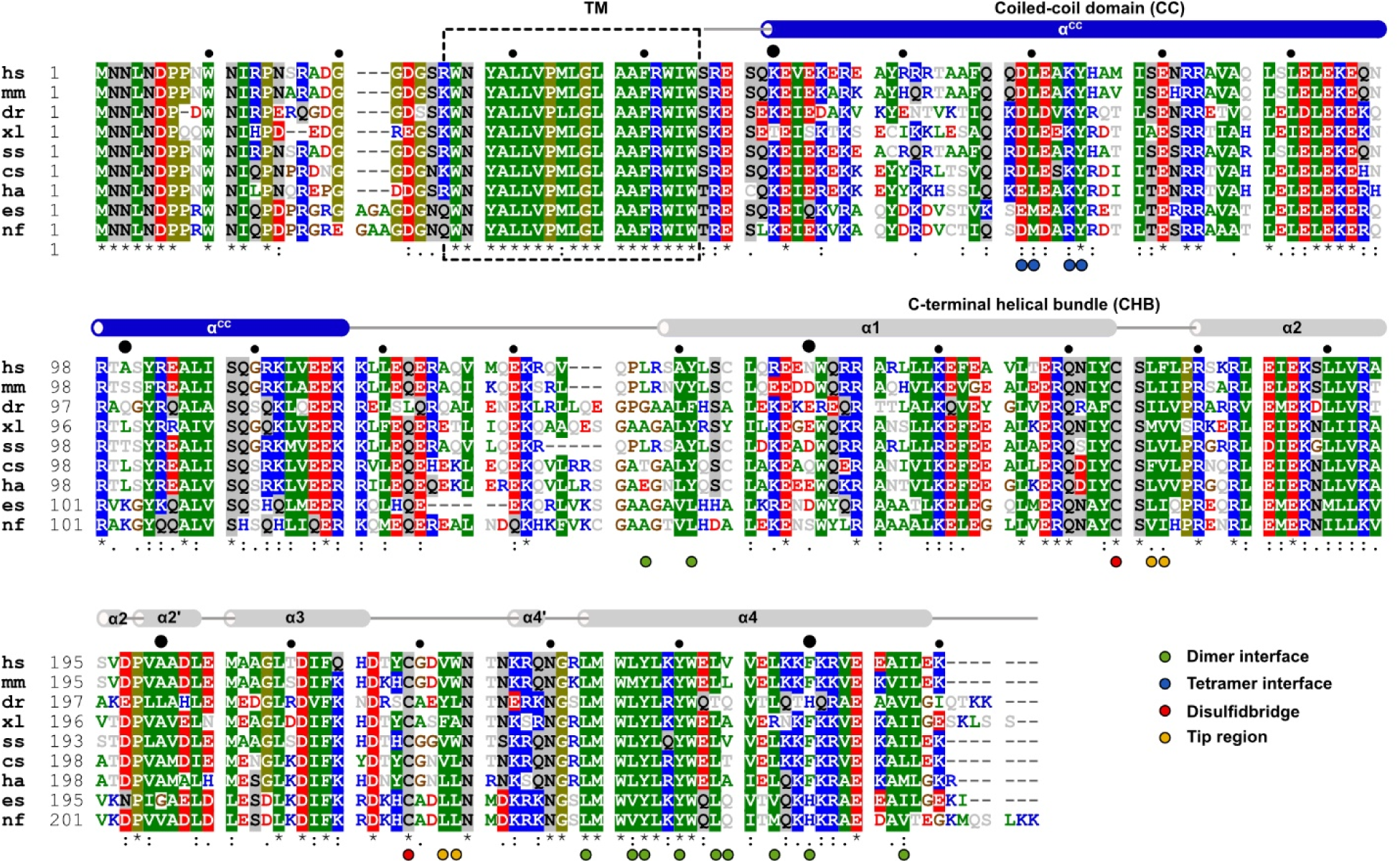
Sequence alignment of CCDC127. **A:** The following sequences are aligned: *Homo sapiens* (hsCCDC127, Uniprot accession code Q96BQ5), *Mus musculus* (mmCCDC127, Q3TC33), *Danio rerio* (drCCDC127, Q1LVA0), *Xenopus laevis* (xlCCDC127, A0A1L8FX28), *Sus scrofa* (ssCCDC127, P0C267), *Chelydra serpentina* (csCCDC127, A0A8C3XK02), *Haliaeetus albicilla* (haCCDC127, A0A7K7NSN9), *Etheostoma spectabile* (esCCDC127, A0A5J5DDH7), *Nothobranchius furzeri* (nfCCDC127, A0A1A8A906). Amino acids are colored according to their chemical and physical properties (positive charge: blue, negative charge: red, hydrophobic: green, proline and glycine: brown, all others: grey). For sequence conservation greater than 70%, the background is highlighted. Residues involved in dimerization, tetramerization, disulfide bond formation and hydrophobic patch formation are labeled with ●, ●, ● and ●, respectively. Small black dots ● represent each 10^th^ and bigger black dots ● each 50^th^ amino acid of human CCDC127. The secondary structure elements obtained from either the crystal structure or the Alphafold3 prediction are show above the sequences.

**Figure S5:**
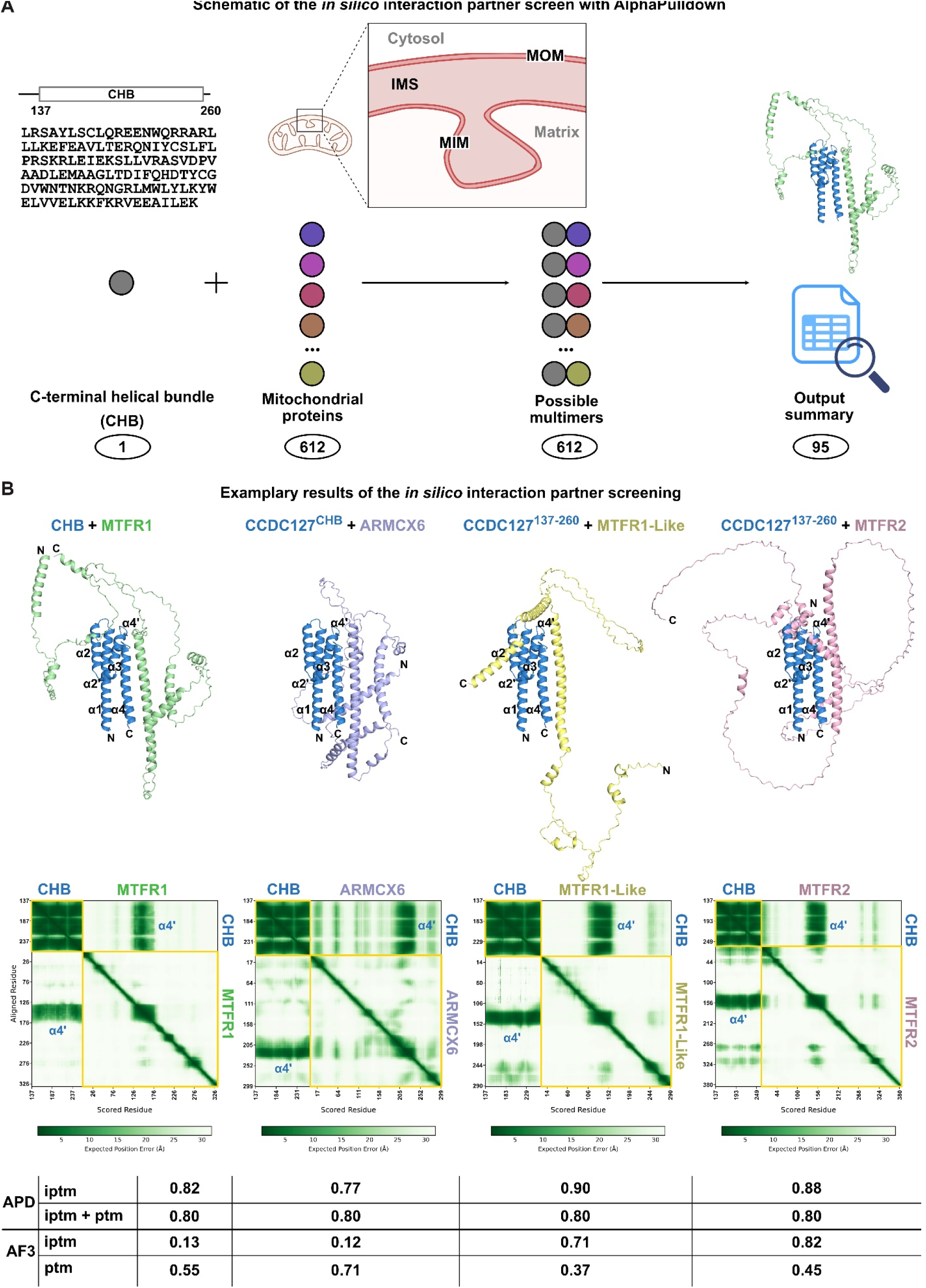
Interaction partners of the CHB identified by AlphaPulldown. **A.** Schematic representation of the AlphaPulldown workflow. The CHB (residues 137-260) of CCDC127 has been used as bait protein, whereas all mitochondrial proteins located in either the mitochondrial outer membrane (MOM), the mitochondrial intermembrane space (IMS) or the mitochondrial inner membrane (MIM) have been used as potential interaction partners (612 proteins). The complex of the CHB with each of the selected mitochondrial proteins has been filter using a threshold of inter-chain PAE of 5, ranked based on the iptm + ptm score and listed in an output sheet (95 proteins). The top hits can be found in **Table S3**. **B.** Structure prediction, PAE plots (Predicted Aligned Error) and respective iptm, ptm and iptm + ptm scores from AlphaPulldown (APD) and Alphafold3 (AF3) runs of selected top hits: MTFR1, Uniprot accession code Q15390; ARMCX6: Uniprot accession code Q7L4S7), MTFR1-like, Uniprot accession code Q9H019; MTFR2, Uniprot accession code Q6P444. All structures are shown as carton representation. The CHB domain of CCDC127 (CCDC127^137-260^) is colored grey, the other proteins differently. The CHB helix α4, which is engaged in the interaction with the other proteins is highlighted in the PAE-plot.

**Figure S6:**
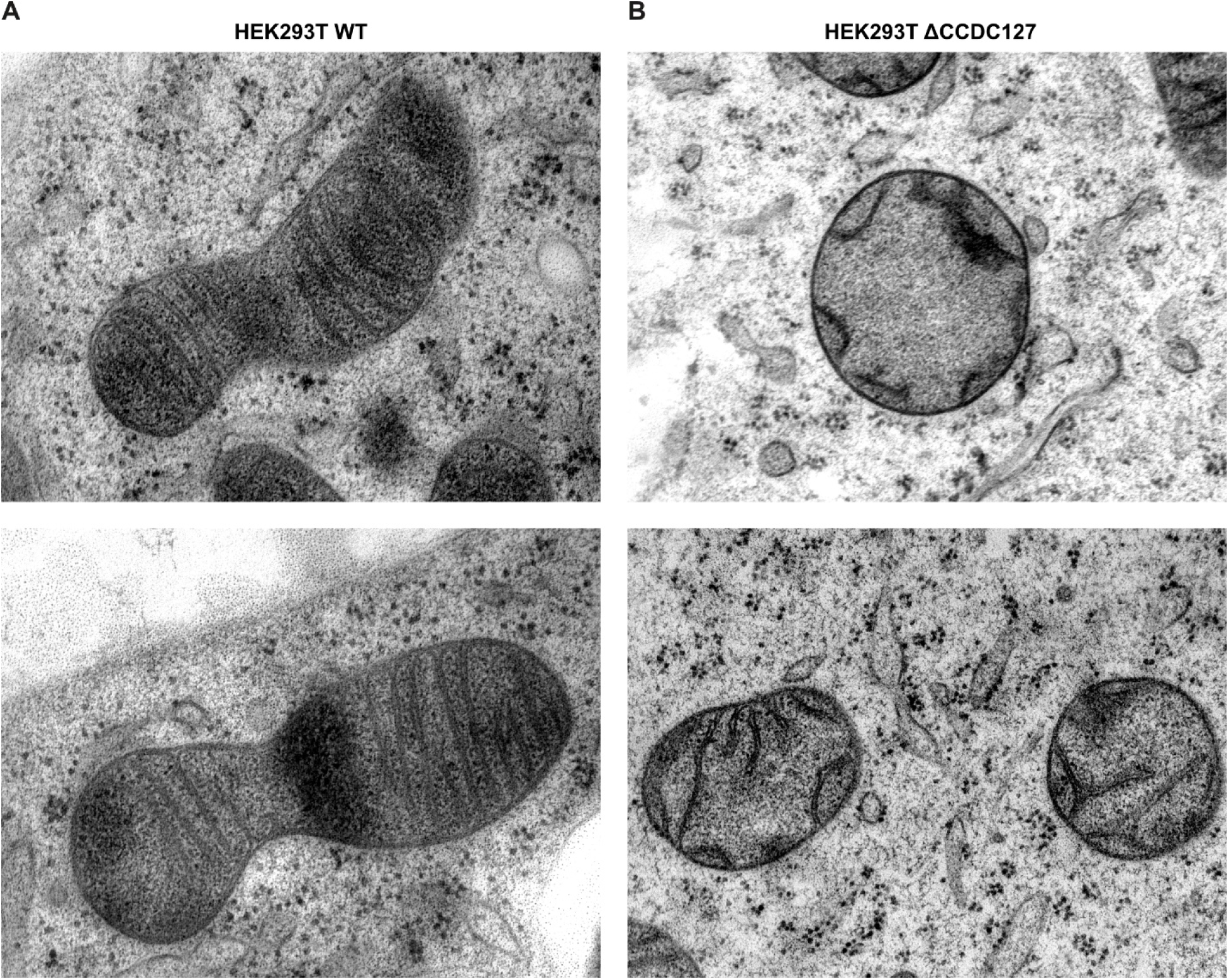
Loss of CCDC127 leads to alterations of mitochondrial membrane architecture. Representative transmission electron microcopy images showing the mitochondrial ultrastructure of wild-type (WT) HEK293T (**A**) and CCDC127 knockout (ΔCCDC127) (**B**) cells. CCDC127-KO mitochondria exhibit shorter and irregularly shaped cristae.

**Table S1:**
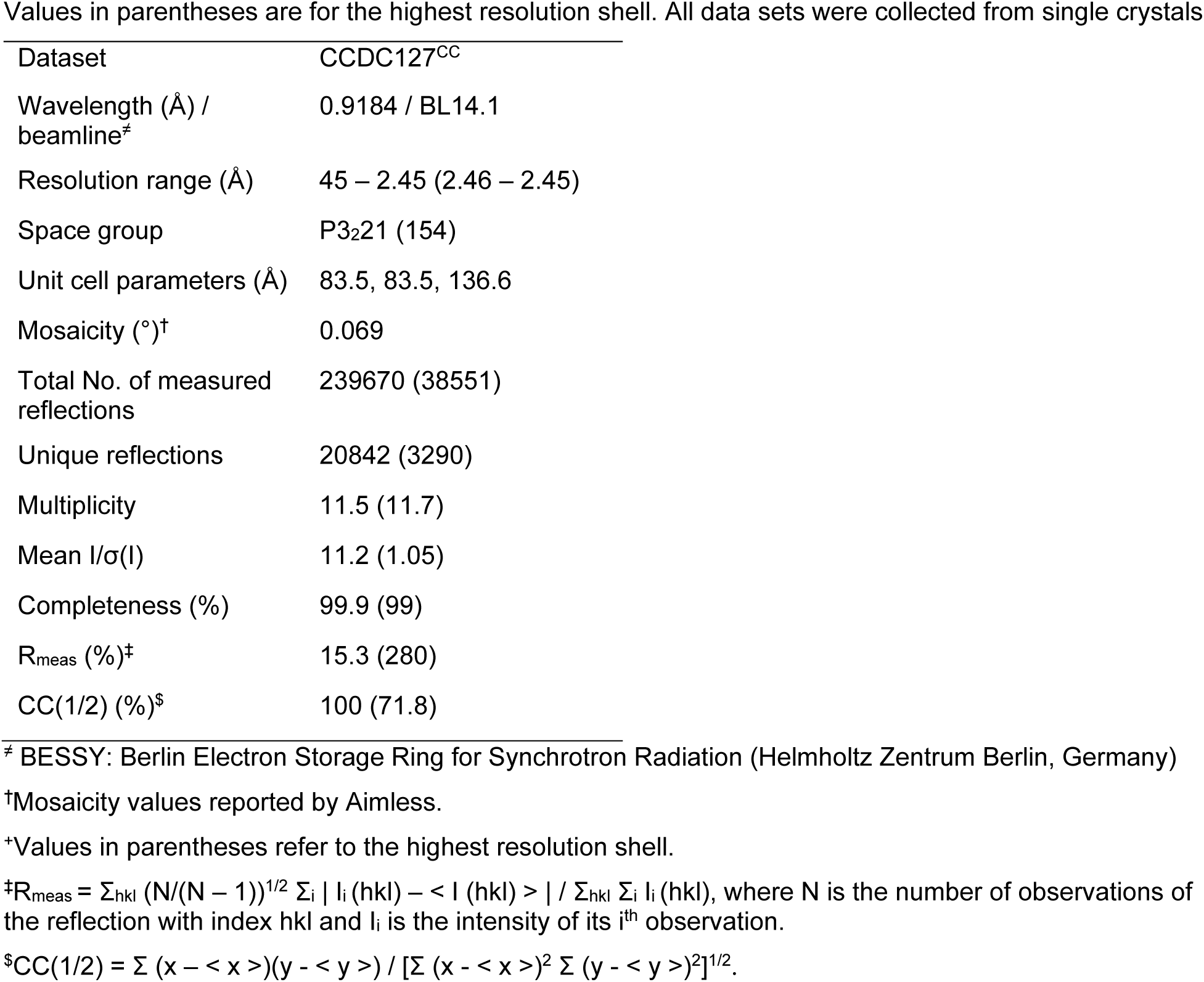
Crystallographic data collection table

Values in parentheses are for the highest resolution shell. All data sets were collected from single crystals
|  |  |
| --- | --- |
| Dataset | CCDC127<sup>CC</sup> |
| Wavelength (Å) /<br>beamline<sup>‡</sup> | 0.9184 / BL14.1 |
| Resolution range (Å) | 45 – 2.45 (2.46 – 2.45) |
| Space group | P3<sub>2</sub>21 (154) |
| Unit cell parameters (Å) | 83.5, 83.5, 136.6 |
| Mosaicity (°)<sup>†</sup> | 0.069 |
| Total No. of measured<br>reflections | 239670 (38551) |
| Unique reflections | 20842 (3290) |
| Multiplicity | 11.5 (11.7) |
| Mean I/σ(I) | 11.2 (1.05) |
| Completeness (%) | 99.9 (99) |
| R<sub>meas</sub> (%)<sup>‡</sup> | 15.3 (280) |
| CC(1/2) (%)<sup>\$</sup> | 100 (71.8) |

<sup>‡</sup> BESSY: Berlin Electron Storage Ring for Synchrotron Radiation (Helmholtz Zentrum Berlin, Germany)
<sup>†</sup>Mosaicity values reported by Aimless.
<sup>\*</sup>Values in parentheses refer to the highest resolution shell.
<sup>‡</sup> $R_{\text{meas}} = \sum_{\text{hkl}} (N/(N - 1))^{1/2} \sum_i |I_i(\text{hkl}) - \langle I(\text{hkl}) \rangle| / \sum_{\text{hkl}} \sum_i I_i(\text{hkl})$ , where N is the number of observations of the reflection with index hkl and I<sub>i</sub> is the intensity of its i<sup>th</sup> observation.
<sup>\$</sup> $\text{CC}(1/2) = \sum (x - \langle x \rangle)(y - \langle y \rangle) / [\sum (x - \langle x \rangle)^2 \sum (y - \langle y \rangle)^2]^{1/2}$ .

**Table S2:** Crystallographic refinement statistics table

Values in parentheses are for the highest resolution shell.
|  |  |
| --- | --- |
| Dataset | CCDC127<sup>CC</sup> |
| Resolution range (Å) | 38 – 2.45 (2.54 – 2.45) |
| R<sub>cryst</sub> (%) | 31.4 |
| R<sub>free</sub> (%) | 34.5 |
| No. of non-H atoms |  |
| Protein | 2656 |
| Ion | 5 |
| Ligand | - |
| Water | 2 |
| R.m.s. deviations |  |
| Bonds (Å) | 0.013 |
| Angles (°) | 1.01 |
| Average B factors (Å<sup>2</sup>) |  |
| Protein | 92 |
| Ion | 92 |
| Ligand | - |
| Water | 78 |
| Ramachandran plot |  |
| Favoured regions (%) | 100 |
| Outliers (%) | 0.0 |
| MolProbity score<sup>#</sup> | 1.69 |
| PDB entry code | 9REU |

<sup>#</sup>As reported by MolProbity at http://molprobity.biochem.duke.edu/

**Table S3:** Top hits of the Alphapulldown *in silico* interaction partner search

| Complex of CCDC127(137-260) with |  | hit # | iptm_ptm | iptm | pDockQ/mpDockQ | AF3 cross validation |  |
| --- | --- | --- | --- | --- | --- | --- | --- |
| Protein name | Genbank accession |  |  |  |  | iptm | pTM |
| CCDC127 | NP_660308 | 1 | 0,86 | 0,90 | 0,47 | 0,82 | 0,85 |
| Bcl-2-like protein 13 isoform i | NP_001257664 | 2 | 0,80 | 0,82 | 0,22 | 0,13 | 0,55 |
| Cytochrome c oxidase subunit I | YP_003024028 | 3 | 0,80 | 0,77 | 0,12 | 0,12 | 0,71 |
| Mitochondrial fission regulator 1 | NP_055452 | 4 | 0,80 | 0,90 | 0,42 | 0,71 | 0,37 |
| ARMCX6 | NP_001171697 | 5 | 0,80 | 0,88 | 0,18 | 0,82 | 0,45 |
| Mitochondrial fission regulator 1-like | NP_001093097 | 6 | 0,80 | 0,88 | 0,33 | 0,88 | 0,42 |
| Mitochondrial fission regulator 2 | NP_001092756 | 7 | 0,78 | 0,89 | 0,38 | 0,87 | 0,39 |
| DNAJ homolog subfamily C member 30 | NP_115693 | 8 | 0,78 | 0,82 | 0,24 | 0,57 | 0,51 |
| Phosphatidate cytidyltransferase | NP_001271330 | 9 | 0,78 | 0,77 | 0,15 | 0,24 | 0,51 |
| Calcium uptake protein 1 | NP_006068 | 10 | 0,78 | 0,81 | 0,12 | 0,21 | 0,48 |
| Peroxisomal carnitine O-octanoyltransferase | NP_001137407 | 11 | 0,78 | 0,75 | 0,09 | 0,15 | 0,79 |
| Bcl-2-like protein 10 | NP_065129 | 12 | 0,78 | 0,78 | 0,15 | 0,54 | 0,66 |
| Cholesterol side-chain cleavage enzyme | NP_001093243 | 13 | 0,78 | 0,75 | 0,07 | 0,13 | 0,62 |
| Bcl-2-like protein 11 | NP_001191039 | 14 | 0,77 | 0,80 | 0,26 | 0,55 | 0,43 |
| Steroidogenic acute regulatory protein | NP_000340 | 15 | 0,77 | 0,77 | 0,40 | 0,63 | 0,63 |
| Stomatin isoform a | NP_004090 | 16 | 0,76 | 0,80 | 0,33 | 0,17 | 0,4 |
| Septin-4 | NP_001185642 | 17 | 0,76 | 0,82 | 0,28 | 0,65 | 0,45 |
| Regulator of microtubule dynamics protein 3 | NP_060615 | 18 | 0,76 | 0,83 | 0,44 | 0,75 | 0,42 |
| ATP synthase peripheral stalk subunit F6 | NP_001676 | 19 | 0,75 | 0,77 | 0,17 | 0,18 | 0,53 |
| Cytochrome c oxidase assembly factor 6 homolog | NP_001193570 | 20 | 0,75 | 0,73 | 0,07 | 0,15 | 0,48 |
| Prohibitin-2 | NP_001138303 | 21 | 0,75 | 0,80 | 0,42 | 0,54 | 0,54 |
| Prohibitin-1 | NP_002625 | 22 | 0,75 | 0,80 | 0,43 | 0,38 | 0,37 |
| Mitochondrial dynamics protein MID49 | NP_631901 | 23 | 0,75 | 0,80 | 0,38 | 0,52 | 0,54 |
| Methylmalonate-semialdehyde dehydrogenase | NP_005580 | 24 | 0,74 | 0,74 | 0,28 | 0,21 | 0,68 |
| Protein PET117 homolog | NP_001158283 | 25 | 0,74 | 0,73 | 0,32 | 0,2 | 0,57 |
| Mitochondrial calcium uniporter regulator 1 | NP_001026883 | 26 | 0,74 | 0,81 | 0,31 | 0,3 | 0,3 |
| Phorbol-12-myristate-13-acetate-induced protein 1 | NP_066950 | 27 | 0,73 | 0,72 | 0,12 | 0,46 | 0,7 |
| Small ribosomal subunit protein mS37 | NP_976043 | 28 | 0,71 | 0,72 | 0,18 | 0,66 | 0,63 |
| S-adenosylhomocysteine hydrolase-like protein 1 | NP_001229603 | 29 | 0,71 | 0,68 | 0,18 | 0,24 | 0,4 |
| MICOS complex subunit MIC60 | NP_001093640 | 30 | 0,70 | 0,79 | 0,15 | 0,34 | 0,28 |
| Coenzyme Q-binding protein COQ10 homolog A | NP_653177 | 31 | 0,70 | 0,71 | 0,15 | 0,17 | 0,47 |
| Coiled-coil domain-containing protein 90B | NP_001182 | 32 | 0,70 | 0,72 | 0,25 | 0,19 | 0,47 |
| Syntaxin-17 | NP_060389 | 33 | 0,70 | 0,73 | 0,18 | 0,31 | 0,39 |

